# Genetic engineering of *Staphylococcus haemolyticus*: Overcoming Restriction-Modification Barriers and Targeting Virulence Genes

**DOI:** 10.64898/2026.02.03.702568

**Authors:** Hermoine J. Venter, J. Pauline Cavanagh, Runa Wolden, Dakota Jones, Richard J. Roberts, Martin O. Christensen, Martha Zepeda-Rivera, Christopher D. Johnston

**Affiliations:** Department of Clinical Medicine, Research Group for Child and Adolescent Health, UiT-The Arctic University of Norway, Tromsø, Norway; Vaccine and Infectious Disease Division, Fred Hutchinson Cancer Center, Seattle, WA 98109; New England Biolabs, Ipswich, MA, 01938-2723, USA; Genomic Medicine, UT MD Anderson Cancer Center, South Campus Research Building, Houston, TX 77054

**Keywords:** *Staphylococcus haemolyticus*, Restriction modification systems, bacterial defence systems, DNA methylation

## Abstract

*Staphylococcus haemolyticus* is an emerging multidrug-resistant nosocomial pathogen noted for robust biofilm formation and complex restriction-modification (RM) systems that hinder genetic manipulation. These barriers have severely limited mechanistic studies into its pathogenesis and immune evasion. Here, we report the development of a molecular toolbox that enables precise genomic engineering of clinical *S. haemolyticus* isolates. Using PacBio Single Molecule Real Time (SMRT) and bisulfite sequencing, we defined the complete genomes and methylomes of nine isolates, generating a functional readout of the active RM defences present in each strain. Among the RM systems identified, a Type II (PDLC03279) and a Type III (PDL3649/PDLC03643) system were significantly overrepresented in clinical isolates, suggesting a potential role in adaptation to host or hospital-associated environments. To bypass these RM barriers, we implemented a dual strategy: first, applying SyngenicDNA-based approaches to eliminate RM target motifs from genetic tools; and second, engineering a surrogate *E. coli* strain (JMC4) to mimic conserved *S. haemolyticus* methylation patterns. These tools significantly enhanced transformation efficiency and enabled targeted knockout of four putative virulence genes (*sraP, secA2, capA*, and *capI*), as well as allelic exchange of the native capsule operon with the corresponding region from a non-encapsulated isolate. To our knowledge, this is the first report of precise genomic modifications in *S. haemolyticus*. The establishment of robust molecular tools for transformation and genome editing lays a foundation for future functional studies of virulence and host adaptation in this resilient opportunistic pathogen.

**Data summary:** All genomic data were deposited in the European Nucleotide Archive and are available in NCBI under the project accession number PRJEB2705. Supporting data are provided in supplementary data files. Accession numbers are given in Table 1.

**Impact statement:** *Staphylococcus haemolyticus* is a multidrug-resistant opportunistic pathogen which causes disease in vulnerable patients. Previously, genetic modification of this pathogen was prevented due to the prevalence of strain-specific restriction modification systems, which use DNA methylation to identify and foreign DNA (including plasmids) when it enters the cells, and then destroy it using restriction enzymes. We analysed the methylomes of a selection of *S. haemolyticus* strains and used this data to tailor molecular tools to evade the RM systems. This enabled us to improve transformation efficiency and perform genome editing, including large-scale chromosomal modifications and targeted deletion of suspected virulence genes (a first for *S. haemolyticus*). Beyond its immediate relevance to researchers studying this pathogen, the tools and approaches developed here have broader utility for genetic engineering of other coagulase-negative staphylococci and bacteria with similar RM barriers. Additionally, we found that two RM systems were enriched in clinical strains, suggesting they may have a function in virulence, in addition to their roles as bacterial phage defence systems.

## Introduction

*Staphylococcus haemolyticus* is a coagulase-negative *staphylococcus* (CoNS) and a typical member of the human skin microbiota, but has also emerged as a significant nosocomial pathogen, particularly in immunocompromised and paediatric patients (1-4). Though historically overshadowed by *Staphylococcus aureus* in clinical settings, *S. haemolyticus* is now the second most frequently isolated CoNS from human blood cultures and is associated with severe hospital-acquired infections, including bacteremia,endocarditis, and sepsis, especially in neonatal intensive care units (5-7). Its clinical importance is further underscored by its high levels of antimicrobial resistance, including multidrug resistance (MDR), and its ability to form robust biofilms on medical devices, which complicate treatment and eradication strategies particularly among at-risk patients (8, 9).

**Table 1.**
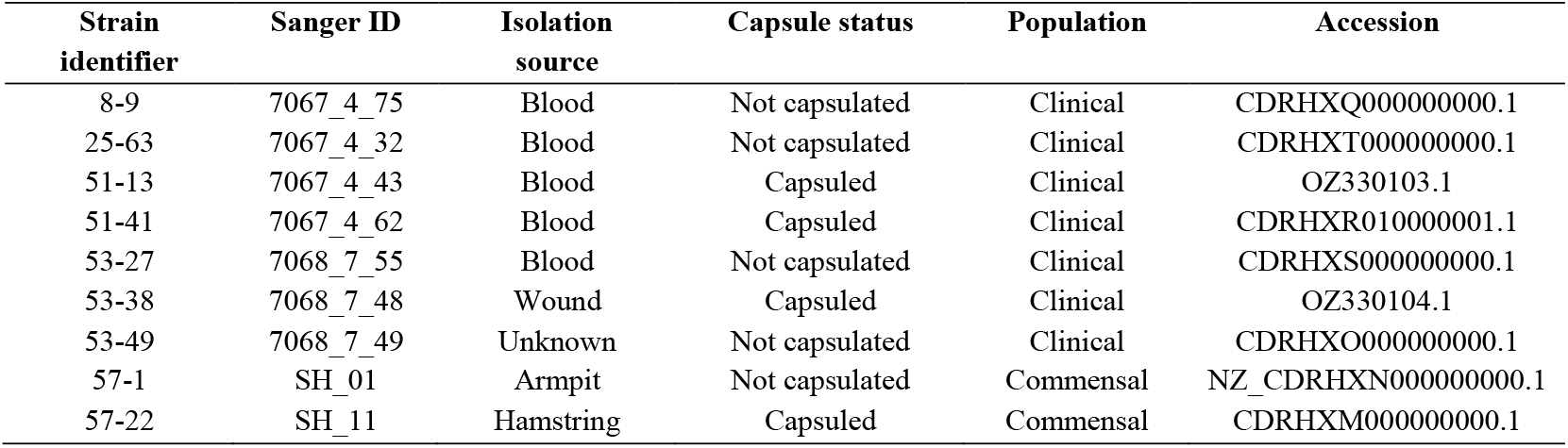
*S. haemolyticus* isolates used in this study.

Despite the growing clinical impact of *S. haemolyticus* and the availability of over 800 sequenced isolates (10), functional studies of its mechanisms of pathogenicity and immune evasion strategies remain scarce (3). This is due in part to the difficulty of genetically manipulating clinical isolates of *Staphylococcus* species, which often harbour complex and strain-specific restriction-modification (RM) systems. These systems protect against foreign DNA via methylation-based self-recognition, degrading unmethylated or improperly modified sequences, mechanisms that evolved to counter phage infection but also block artificial transformation with *in vitro*-constructed plasmids (11). While similar barriers in *S. aureus* have been overcome with advanced genetic tools (12-15), no such systems existed for *S. haemolyticus*, limiting mechanistic investigation of its virulence strategies.

Due to the hypervariability and strain specificity of RM system target motifs (11, 16-19), overcoming these defences requires detailed knowledge of the specific systems present in each clinical isolate. This information can be inferred from the bacterial methylome, the complete set of DNA methylation marks across a genome. Methylation-sensitive sequencing methods include PacBio Single-Molecule Real-Time (SMRT) sequencing (20), Nanopore sequencing (21), bisulfite sequencing (22) or Enzymatic Methyl-seq (23) approaches. Once an isolate’s methylome is defined, two primary approaches can be used to evade RM systems: mimicry-by-methylation and RM-silencing (12). These approaches modify DNA sequences in different ways but are both tailored to prevent destruction by restriction enzymes (REase) present in the strain of interest and increase transformation efficiencies.

Mimicry-by-methylation (also known as plasmid-artificial-modification) involves passaging genetic tools through an intermediate host engineered to replicate the target strain’s methylation profile (13), thereby protecting the DNA from degradation upon transformation (13-15). In contrast, RM-silencing, (also known as SyngenicDNA-based approaches), uses methylome information to redesign the DNA sequence of genetic tools by removing unnecessary (non-functional or superfluous) regions that harbour RM target motifs, or by introducing synonymous codon substitutions that eliminate specific RM target motifs. These sequence-adapted ‘RM-silent’ tools are synthesized de novo and evade recognition and cleavage by individual RM systems upon transformation (12, 24).

To enable functional genetic analysis of *S. haemolyticus*, we applied both mimicry-by-methylation and SyngenicDNA-based strategies to establish a comprehensive genetic system for engineering clinical isolates. We first defined the complete genomes and methylomes of nine *S. haemolyticus* isolates (seven clinical, two commensal), enabling the design of RM-silent tools. Using these tools, we optimised competent cell and transformation protocols across multiple *S. haemolyticus* strains. Further, we constructed an intermediate cloning host (*Escherichia coli* JMC4) to support plasmid-artificial modification approaches for specific isolates which harboured a conserved *S. haemolyticus* RM system identified in this work.

To demonstrate the utility of this platform, we performed extensive allelic exchange in a clinical *S. haemolyticus* isolate, generating a series of targeted isogenic mutants with chromosomal alterations ranging in size from 600 bp - 17 Kbp. As a proof of concept, we targeted several putative virulence genes. For example, the <u>S</u>erine <u>R</u>ich <u>A</u>dhesin for <u>P</u>latelets gene (*sraP*), more prevalent in clinical than commensal isolates of *S. haemolyticus* (3), encodes a surface protein putatively involved in host-cell adhesion and biofilm formation, which contributes to pathogenicity in *S. aureus* and other gram positive bacteria (25, 26). We also targeted the related *secA2* gene, encoding an accessory secretory factor, whereby deletion disrupts surface localization of the SraP protein (27). Additionally, as capsule expression is frequently linked to immune evasion and virulence (28), we modified the capsule polysaccharide operon, which we previously showed varies among *S. haemolyticus* clinical isolates and differs from that of the JCSC 1435 type strain (3, 4, 28). By enabling reliable genome editing in clinical isolates, this work overcomes key technical barriers in the mechanistic study of *Staphylococcus haemolyticus* virulence, immune evasion, and strain-level variation.

## Methods

### Isolates

Our collection consists of 169 previously sequenced *S. haemolyticus* strains. Of these, 123/169 are of clinical origin, and 46/169 are commensal isolates from non-symptomatic carriers (PRJEB2705, (3)). Prophage regions and phage defence systems were identified using the PHASTEST (29) and PADLOC (30) webservers. kSNP4 was used to construct a reference-free phylogenetic tree using the whole genome sequences (kmer=17, FCK=0.650) (31). The resulting parsimony tree was visualized and annotated with PADLOC data using iTOL v7 (32). PADLOC data was used to identify and locate restriction-modification systems across genomes. Maximum-likelihood dendrograms were generated based on MUSCLE-aligned amino acid sequences using MEGA v11 (33) and visualized using iTOL v7 (35). Nine *S. haemolyticus* isolates were selected for further analysis. Seven were clinical isolates obtained from patients in hospital settings, and two were commensal isolates collected from asymptomatic, healthy volunteers.

### Methylation analyses

High-molecular-weight genomic DNA was isolated using the MasterPure™ Gram Positive DNA Purification Kit (Lucigen, USA) with modifications (see supplementary methods). DNA was submitted to the Norwegian Sequencing Centre for PacBio Single Molecule Real Time (SMRT) sequencing. Circular Consensus Sequencing (CCS) reads were demultiplexed and assembled with Canu v1.8 (34), circularised with Circlator (35), and polished with Pilon (36). Methylated motifs were identified using SMRT Link Base Modification and Motif Analysis (BMMA) pipeline (v 6.0.0.47841, SMRT Link Analysis Services and GUI v 6.0.0.47836).

As SMRT sequencing has limited sensitivity for 5-methylcytosine (5mC), we supplemented this analysis with bisulfite sequencing (22). Unmethylated cytosines were converted to uracil using the EpiMark Bisulfite Conversion Kit (New England BioLabs), followed by PCR amplification using EpiMark® Hot Start Taq (primers listed in Supplementary Table 1), and sequencing. Cytosine methylation was inferred by comparison with unconverted genomic sequences. Closed genomes and methylation data were submitted to REBASE (37) for RM system annotation.

### Evading RM systems using SyngenicDNA (SynDNA)

RM-silent (SynDNA) versions of the minicircle-forming plasmid pEPSA5MC were constructed as we described previously (12). For isolate 51-13, pEPSA5MC was edited *in silico* to remove G<u>A</u>GG/<u>C</u>CTC (Type II) RM target motifs. The modified construct was synthesized in three parts by Synbio Technologies (Monmouth Junction, USA) and assembled using NEBuilder Hi-Fi Assembly Master Mix (New England BioLabs, USA). The assembled plasmid was transformed into the minicircle-producing *E. coli* strain JMC1 (12). Minicircle production was induced with arabinose, and minicircle DNA was isolated using the NucleoSpin kit prior to transformation into *S. haemolyticus* (see Supplementary Methods). SynDNA minicircles were used to optimize transformation efficiency by adjusting electroporation and recovery conditions (38).

### Evading RM systems using Mimicry-by-Methylation

Fifty-three isolates in our *S. haemolyticus* collection harboured a Type II RM system (MTase PLDC03279) that targets the GAGG/CCTC motif. This system was both highly prevalent and difficult to circumvent via SynDNA sequence editing, as GAGG/CCTC motifs occur frequently and existed in regulatory regions of the plasmid. To enable plasmid transformation in strains carrying this RM system, we constructed a surrogate *E. coli* strain capable of replicating G^m6^AGG/^m4^CCTC methyl-modification on passaged plasmids (see Supplementary Methods).

Using a recombineering strategy (39), we integrated the *S. haemolyticus* MTase PDLC03279 gene into *E. coli* strain ZYCY10P3S2T while simultaneously deleting the native *dcm* gene, which encodes an *E. coli* MTase associated with C^m5^CW<u>G</u>G modification often targeted by Type-IV restriction systems in *Staphylococcus* species. Integration was confirmed by PCR and whole-genome sequencing. Successful methylation of the GAGG/CCTC motif was validated by digestion of plasmid DNA with the restriction enzyme MnlI, and further by methylome analysis using SMRT Link Base Modification and Motif Analysis (BMMA) pipeline (v 6.0.0.47841, SMRT Link Analysis Services and GUI v 6.0.0.47836). The resulting strain, designated JMC4, supports methylation mimicry of the enriched *S. haemolyticus* RM target motif (G^m6^AGG/^m4^CCTC) and produces minicircle DNA constructs for downstream transformation.

### Allelic exchange

Isolate 53-38 was selected for genetic manipulation based on: i) its clinical relevance, ii) its uncharacterized but prevalent capsule type (3), iii) the presence of *sraP* and the accessory sec transport system, as well as iv) its RM system being representative of many other *S. haemolyticus* strains within our collection. The targeted genes and operons are listed in Table 2.

**Table 2.** Genes and regions modified in this study.

| Name | Gene product(s) | Size (bp) | Type of modification |
| --- | --- | --- | --- |
| *sraP* | Serine-rich adhesin for platelets | 9600 | Knockout |
| *secA2* | Protein translocase subunit | 2931 | Knockout |
| *capA* | Tyrosine-protein kinase<br>transmembrane modulator EpsC | 663 | Knockout |
| *capI* | Glycosyltransferase | 1113 | Knockout |
| Capsule operon | Proteins involved in capsule<br>synthesis and transport | 17 090 | Knockout/Knock-in |
| *EpSN* | DegT family Aminotransferase | 231 | Knock-in |
| *tagU* | LCP family protein | 948 | Knock-in |

Allelic exchange was performed using plasmid pIMAY-Z (a gift from Dr Clement Ajayi) which carries a chloramphenicol resistance cassette and the *lacZ* operon for phenotypic screening of plasmid integration and excision. Plasmids were assembled using the protocol of Monk and Stinear (14), except using NEBuilder HiFi (in place of SLiCE), and transformation conditions were adapted for *S. haemolyticus*. For each gene, flanking regions (∼500 bp upstream and downstream) were amplified from isolate 53-38 genomic DNA using Q5® High-Fidelity 2X Master Mix (New England BioLabs). The plasmid backbone was linearized by PCR, and recombination templates were assembled into pIMAY-Z. Constructs were transformed into *E. coli* NEB 5-alpha, screened by colony PCR, and sequence-verified. When needed, Type I RM target motifs were removed via site-directed mutagenesis before passaging plasmids through JMC4. Because direct mutants were not obtained, we followed the slow integration protocol (Monk 2021, Monk 2013; see supplementary methods). Following plasmid excision, colony PCR was used to identify knockouts. Positive clones were confirmed by whole-genome sequencing (Azenta).

## Results

### *S. haemolyticus* isolates harbour strain-specific and conserved restriction-modification systems

Methylated motifs identified by PacBio SMRT and bisulphite sequencing are summarized in Table 3 (see also Supplementary Table S4). Genome analysis confirmed that the majority of isolates in our full collection (156/169, 92.6%) encode at least one methyltransferase (MTase), and DNA methylation was detected in all but one of the nine isolates analysed by SMRT-sequencing. This is consistent with estimates that ∼93% of bacterial species possess DNA methylation genes (18, 40, 41). Methylation-lacking isolates were disproportionately commensal: only 3 of 123 (2.4%) clinical isolates lacked MTases, compared to 10 of 46 (21.7%) commensals (p<0.0001, Chi-square test with Yates’ correction). Despite this, most carried alternative phage defence systems, including Type IV RM systems, CBASS systems, CRISPR loci and other phage defence genes (see Figure 2 for the distribution of phage defence systems by strain, and supplementary Figure S1 for the number of strains that contained each defence system). These systems were often co-located in “defence islands,” a hallmark of horizontally acquired immune regions in prokaryotes (42, 43). Several MTase genes were identified within prophage regions, consistent with the reported role of phage-encoded MTases in protecting phage DNA from host endonucleases during infection (44, 45).

**Table 3.** RM systems and methylation motifs identified in *S. haemolyticus* isolates by PacBio SMRT sequencing and bisulphite sequencing, along with the distribution of these RM systems among clinical and commensal isolates. RM systems were predicted using PADLOC (30) and REBASE (37). Methyl-modifications were assigned to specific RM systems via REBASE analysis (see Supplementary Table S4). Orphan methylase lack a detected cognate restriction enzyme. MTase = methyltransferase, REase = restriction enzyme, Specificity = specificity subunit.

| System | PADLOC accession number | Function | Methyl-modified motif | Clinical (n-123) | Commensal (n-46) | Total |
| --- | --- | --- | --- | --- | --- | --- |
| TI | PDLC03040 | REase |  | 16 | 5 | 21 |
|  | PDLC03019 | MTase |  | 15 | 3 | 18 |
|  | PDLC03068 | Specificity | CRTANNNNNNTTC | 6 | 0 | 6 |
|  | PDLC03164 | Specificity | TTACNNNNNTAC | 8 | 3 | 11 |
|  | PDLC03047 | REase |  | 11 | 0 | 11 |
|  | PDLC03021 | MTase |  | 11 | 0 | 11 |
|  | PDLC03107 | Specificity | TAANNNNNNRTGG | 11 | 0 | 11 |
|  | PDLC03048 | REase |  | 9 | 9 | 18 |
|  | PDLC03022 | MTase |  | 8 | 8 | 16 |
|  | PDLC03165 | Specificity | RAAGNNNNNNTATT | 8 | 9 | 17 |
| TII | PDLC03514 | REase |  | 51 | 0 | 54 |
|  | PDLC03279 | MTase | GAGG/CCTC | 50 | 3 | 53 |
|  | PDLC03286 | MTase | Unknown. Orphan | 4 | 0 | 4 |
|  | PDLC03294 | MTase | Unknown. Orphan | 35 | 2 | 37 |
|  | PDLC03553 | REase |  | 17 | 8 | 25 |
|  | PDLC03329 | MTase | GATC | 17 | 8 | 25 |
|  | PDLC03726 | MTase |  | 9 | 0 | 9 |
| TIII | PDLC03649 | REase |  | 55 | 3 | 58 |
|  | PDLC03643 | MTase | GGAG | 54 | 2 | 56 |
|  | PDLC03641 | MTase | VGACAT | 1 | 1 | 2 |
| TIV | PDLC03658 | mREase |  | 54 | 11 | 65 |

**Figure 1.**
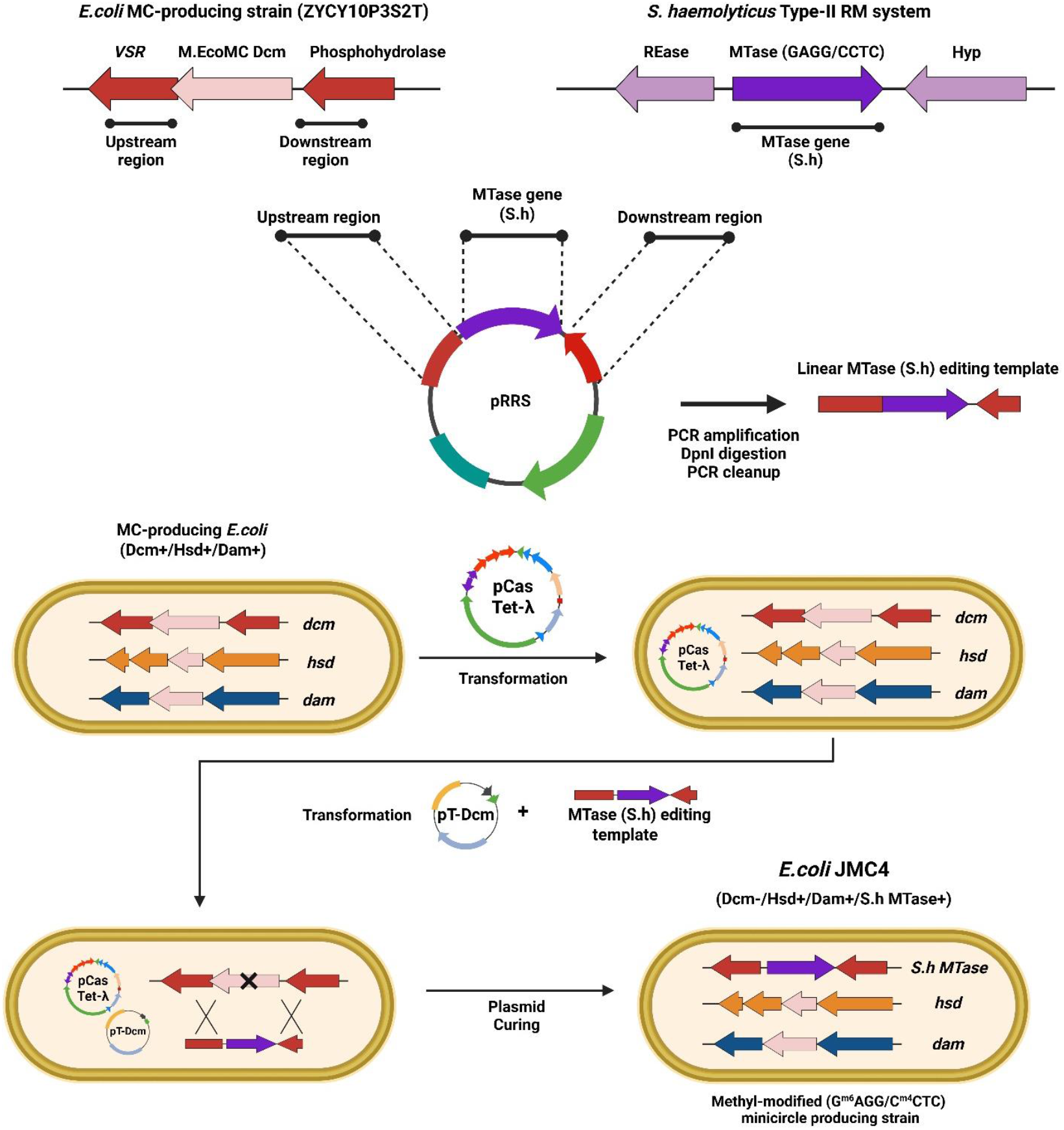
Construction of JMC4, a minicircle-forming *E. coli* which replicates the G**A**GG/**C**CTC methylation pattern found in *S. haemolyticus*.

**Figure 2.**
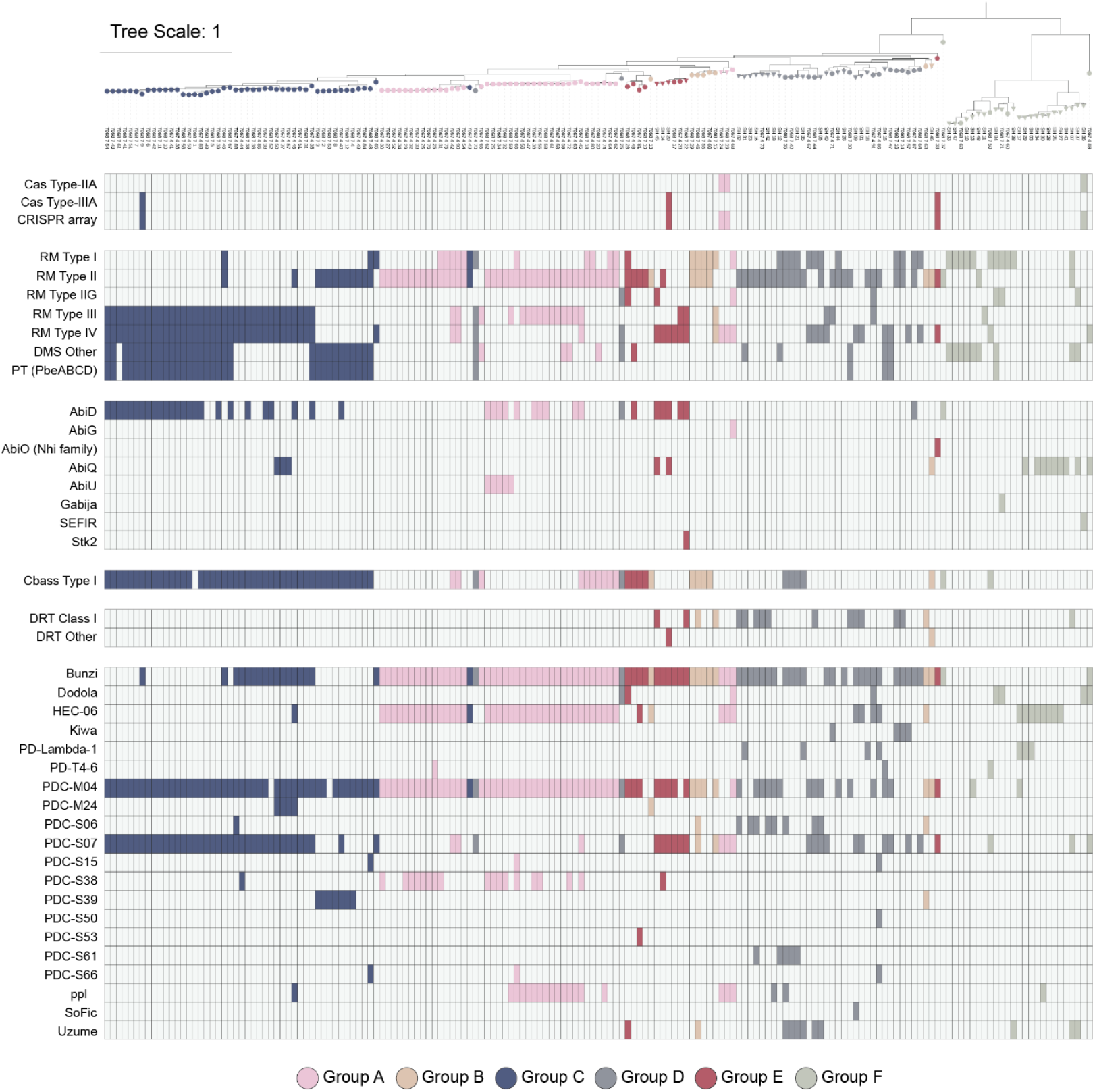
A reference-free phylogenetic tree constructed from whole genome sequences showing the distribution of phage defence systems (as identified by PADLOC) across our collection of *S. haemolyticus* genomes. Circles = clinical isolates, triangles = commensal isolates.

Type I systems are made up of three components, referred to as host specificity for DNA (*hsd)* genes: an MTase (HsdM), an REase (HsdR) and a specificity subunit (HsdS). We mapped the presence of the *hsd* genes across the phylogenetic tree for our collection of genomes. Fifty (50/169, 29.58%) isolates contained Type I systems. Like *S. epidermidis*, the Type I systems in *S. haemolyticus* occur in complete operons and are often associated with the hypervariable staphylococcal cassette chromosome (SCC) region of the genome. The dendrograms (supplementary figures S2A-C) show which MTases, REases and specificity subunits consistently occur together. One system (pale blue) typically contained two specificity subunits, which may indicate the possibility of phase variation (46). Unlike *S. aureus*, where specificity subunits tend to cluster by clonal complexes (47), no such tendency to cluster by clade was observed in *S. haemolyticus*.

Type II systems are highly prevalent in *S. haemolyticus*, particularly among clinical strains (supplementary figure S4). In our collection, 115/169 (68.05%) isolates contained a Type II system, with 96/123 clinical isolates and 19/46 commensal isolates (p<0.0001, Chi-square test with Yates’ correction). This included 90/169 (53.25%) isolates that contained Type II systems composed of an MTase and cognate REase, 53/169 (31.36%) which lacked REases (orphan MTases, see DMS other on Figure 2), and 8/169 (4.73%) Type IIG systems composed of bi-functional enzymes possessing both MTase and REase activity. It was not uncommon for isolates to have multiple Type II systems. The dendrogram of the Type II MTases (supplementary figure S3A) shows 18 branches corresponding to PADLOC-identified proteins.

Type II MTase PDLC03279 was notable for methylating both adenine and cytosine within its G^m6^AGG/C^m4^CTC recognition motif, resulting in asymmetric methylation on each strand. PDLC03279 (along with its cognate REase, PDLC03514) was significantly enriched in clinical strains, appearing in 41% of clinical isolates but only 6.5% of commensals (p<0.0001, Chi-square test with Yates’ correction).

The Type III system containing MTase PDLC03643 (recognizing the similar but distinct GG^m6^AG motif) was also significantly enriched among clinical strains, present in 43.9% of clinical isolates versus 4% of commensals (p<0.0001, Chi-square test with Yates’ correction). One other Type III system was present, MTase PDLC0364, which was only present in one commensal isolate, SH11.

One well-conserved Type IV system, PDLC03658, was detected and was present in 65/169 (38.46%) of isolates and found in all clades. This system shares 75.97% similarity with the *S. aureus* Type IV enzyme, SauUSI, which recognises the S5mCNGS motif (48).

While some RM systems were relatively common, most isolates exhibited unique combinations of Type I– III RM systems, orphan MTases, and phage-encoded methylation genes, resulting in isolate-specific methylomes. This strain-specific variability is consistent with observations across other *Staphylococcus* species (12-15, 47, 49).

### Methylome-guided strategies permit evasion of RM systems during engineering of *S. haemolyticus*

We identified that specific Type II and Type III systems were conserved across several isolates of *S. haemolyticus*, suggesting that development of methods to evade these systems could have broad utility across several strains of interest to the field. Thus, we invested our efforts in the construction of an *E. coli* strain (JMC4) that could be used for artificial methyl-modification of shuttle vectors for *S. haemolyticus*. Type II MTase are more readily subject to such recombinant mimicry-by-methylation approaches (as their MTase often exists as a distinct and separate gene from their cognate REase gene), and thus we focused on the PDLC03279 MTase (G^m6^AGG/C^m4^CTC) enriched in clinical isolates.

Whole-genome sequencing and MnII digestion confirmed successful expression of the PDLC03279 MTase and methyl-modification of G^m6^AGG/C^m4^CTC motifs in JMC4 (Figure 3). Additionally, targeted sequence modification that removed RM target motifs further led to an increase in transformants. While the original pEPSA5 plasmids (8361 bp) resulted in no transformants, pEPSA5 minicircles (2668 bp) produced in an earlier mini-circle production strain (JMC1) lacking the PDLC03279 MTase resulted in 2.2 × 10^2^ CFU/µg DNA. Subsequent combined use of the SynDNA edited pEPSA5 minicircles and the JMC4 strain yielded a 6.4 × 10^2^ CFU/µg (a 2.2-fold increase) in the number of transformants compared to unmodified pEPSA5 minicircles (see Figure 3). After further protocol optimization, transformation with pEPSA5 minicircles reached up to 1 × 10^4^ CFU/µg DNA. Transformation efficiency for larger plasmids such as pIMAY-Z (9800 bp) was lower (1 × 10^2^ CFU/µg DNA), but still sufficient for subsequent allelic exchange experiments.

**Figure 3:**
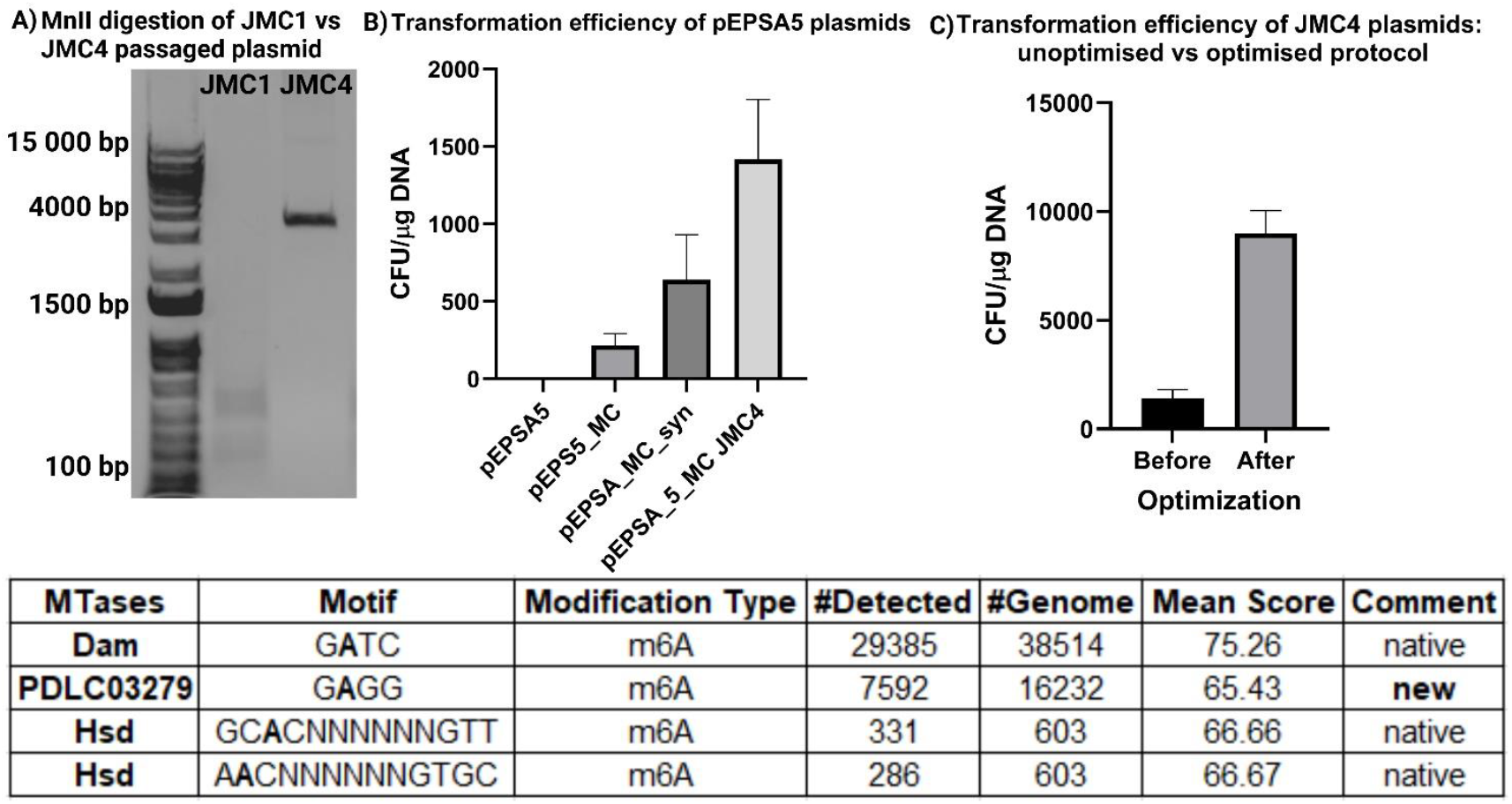
**A**: MnII digestion of pEPSA5 plasmids isolated from JMC1 and JMC4. The recognition sequence of MnII is CTCC. DNA which is methylated at that sequence is protected from digestion by MnII. **B:** Improvements in transformation efficiency of plasmid pEPSA5 and its minicircle form. **C:** Further improvements in transformation efficiency of pEPSA5_MC passaged through JMC4, when the protocol was optimized. The table summarizes the methylated motifs detected in JMC4 following PacBio SMRT sequencing.

### Targeted genome editing in a *S. haemolyticus* clinical isolate

To validate the effectiveness of our RM-silent tools in functional genomic applications, we performed targeted gene deletions and operon replacement in *S. haemolyticus* clinical isolate 53-38. This strain was selected due to its clinical relevance, the presence of a capsule type common to clinical isolates that remains poorly characterized, and the presence of the conserved PDLC03279 RM system.

Using JMC4 and SynDNA-based tools we developed, we successfully knocked out four putative virulence genes, namely *sraP, secA2, capA*, and *capI* (Figure 4, Table 2). These genes were selected based on their putative roles in adhesion (*sraP*), protein export (*secA2*), and capsule biosynthesis (*capA, capI*), which are hypothesized to contribute to *S. haemolyticus* immune evasion and persistence in clinical settings. Targeted deletions were confirmed by PCR screening and validated by whole-genome sequencing, which verified precise allelic exchange without off-target modifications.

**Figure 4.**
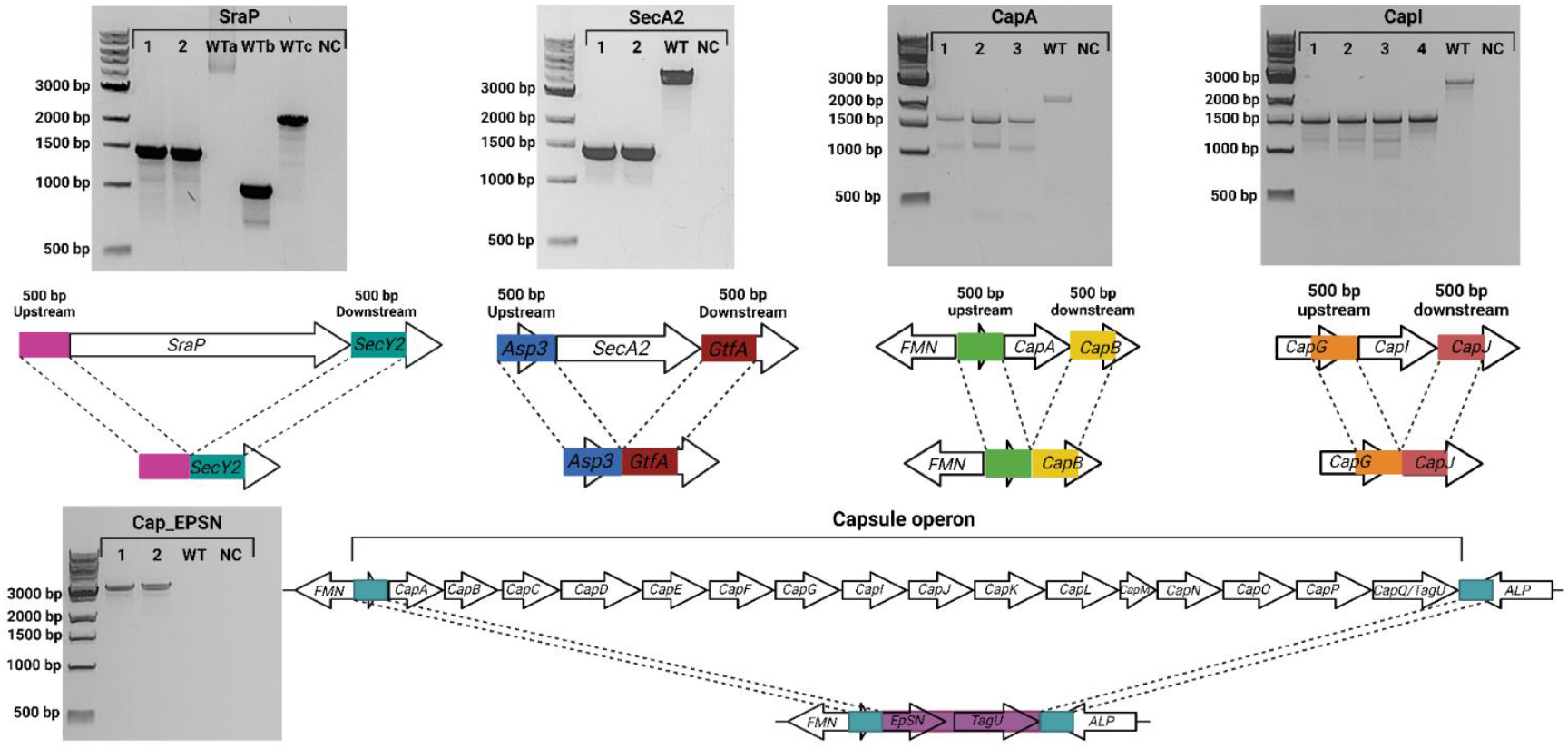
To construct allelic exchange plasmids, the flanking regions (500 bp) from up- and downstream of the gene of interest were cloned into plasmid pIMAY-Z. If Type I target motifs were present, they were removed by site-directed mutagenesis. All plasmids were propagated in E. coli JMC4 to methylate the G^m6^AGG/C^m4^CTC recognition motifs, before electroporation into isolate 53-38.

In addition to generating targeted deletions, it is advantageous to simultaneously replace specific genes with cassettes or marker genes, and to show this is technically feasible using our tools, we swapped the entire capsule operon (∼17 Kbp) of isolate 53-38 with two genes (*EpSN* and *tagU*) that are present in the same genomic locus of the non-encapsulated isolate 53-49 (Figure 4). It is notable that this approach was successful in generating a mutant strain, while previous efforts, which sought to solely eliminate the capsule operon, were not, potentially suggesting essentiality of genes within these regions. Further, this knock-in strategy was designed to mimic a naturally occurring capsule-deficient *S. haemolyticus* genotype, enabling future studies of capsule-associated phenotypes and immune interactions.

## Discussion

*S. haemolyticus* is an opportunistic pathogen that disproportionately affects vulnerable patients (6, 9). Despite its clinical relevance, the lack of molecular tools has historically impeded functional studies of its virulence and immune evasion mechanisms. In this study, we address this gap by constructing a methylation-mimicking *E. coli* strain (JMC4) capable of replicating the methyl-modified G^m6^AGG/C^m4^CTCC motif that is conserved across many *S. haemolyticus* clinical isolates. In parallel, we successfully applied SynDNA-based strategies to eliminate additional RM recognition motifs from the underlying sequence of genetic tools, demonstrating that they retain functionality. Together, these approaches enabled us to overcome barriers of restriction modification and perform efficient transformation and precise genomic modification within *S. haemolyticus* strains.

Using this platform, we successfully generated knockouts of four genes implicated in virulence (*sraP, secA2, capA*, and *capI*) with deletion sizes ranging from 663 bp to 9600 bp. We also performed a large-scale replacement of the 17 Kbp capsule operon with genes from a non-encapsulated isolate, demonstrating this system’s capacity for complex, scarless genome engineering. To our knowledge, this is the first report of targeted chromosomal modifications in any *S. haemolyticus* strain. These tools should provide a robust foundation for a range of more advanced applications, including transposon mutagenesis, CRISPR-Cas9 and CRISPRi gene editing, and integration of reporter constructs (50-52) for *S. haemolyticus*.

Our methylome analysis revealed signatures of phage-host coevolution. A recent paper by da Silva and Rossi (49) analysed 692 *S. haemolyticus* genomes and demonstrated that this species has a diverse inventory of phage defence systems. We have added to this developing view of *S. haemolyticus* and have identified multiple RM systems (including some phage-encoded MTases and anti-restriction genes) as well as numerous other phage defence systems, further highlighting that *S. haemolyticus* is engaged in an ongoing evolutionary arms race with its phage predators (40, 42). Furthermore, our methylome analysis indicates that while many RM systems are active under standard laboratory conditions, some (including those located in phage regions) are seemingly inactive. da Silva and Rossi (49) found that many phage defence genes were not constitutively expressed and inferred that expression may be linked to environmental triggers or stressors.

Two RM systems, PDL3649/PDLC03643 (Type III, GG^m6^AG) and PDLC03279 (Type II, G^m6^AGG/C^m4^CTC), were significantly enriched in clinical isolates compared to commensal strains. This overrepresentation suggests that these systems may confer an adaptive advantage in hospital or host environments, possibly through enhanced phage resistance or epigenetic regulation of gene expression. DNA methylation has been shown to influence phase variation and virulence in other pathogens (53-56), and these systems warrant further investigation as they may play similar roles in *S. haemolyticus*. Further investigation into their regulatory functions and potential involvement in *S. haemolyticus* pathogenicity may yield new insights into host adaptation and identify novel therapeutic targets for combating this nosocomial opportunistic pathogen.

## Conclusion

This study marks a major advance in the genetic engineering of *Staphylococcus haemolyticus*, an emerging multidrug-resistant pathogen with complex, strain-specific restriction-modification systems. By developing SyngenicDNA tools and the methylation-mimicking *E. coli* strain JMC4, we enabled precise genome editing in clinical isolates. The ability to generate targeted deletions and large chromosomal modifications establishes a scalable platform for mechanistic studies and therapeutic target discovery, laying the groundwork for broader genetic tractability across coagulase-negative staphylococci.

