## Supplementry tables S1-3 for "Genetic engineering of *Staphylococcus haemolyticus*: Overcoming Restriction-Modification Barriers and Targeting Virulence Genes"

**Supplementary Table S1) Primers used in this study. Overlaps are underlined and bolded.**

| Primer name | Primer sequence (5’-3’) | Characteristics/References |
|---|---|---|
| **Bisulfite sequencing** | | |
| B1_CON_FW | TGTTTTGAATTATTTAATTATTTTGGT | Amplification of bisulfite-converted DNA |
| B1_CON_RV | ACACTTATAACACTTACACAC | Amplification of bisulfite-converted DNA |
| B2_CON_FW | GTATAATTATTAAAAATGTAAAGTTTGTT | Amplification of bisulfite-converted DNA |
| B2_CON_RV | CTTACAATATTAAATTTCTATAACACACAC | Amplification of bisulfite-converted DNA |
| B3_CON_FW | CACATATTAAAAACAAAAAAAATAAAATATATATC | Amplification of bisulfite-converted DNA |
| B3_CON_RV | GTTTTTATATAAATATTATGTTTTATAATTTTTATTTTAG | Amplification of bisulfite-converted DNA |
| B4_CON_FW | TGGTGTTTTGTGAGGTGATGT | Amplification of bisulfite-converted DNA |
| B4_CON_RV | AAAAAAATCATTAATTACTTAAATACTAAAAATCCTTTTA | Amplification of bisulfite-converted DNA |
| **Syngenic pEPSA5 (GAGG)** | | |
| Syn_PartA_F | **GAGCTATTAAGCCGA**CCATTCGACAAGTTTTG | Overlap |
| Syn_PartA_R | CTGTTAGCACTTATATTTTGTTATTGTTCTTCTTCGATTTC | Overlap |
| Syn_PartB_F | **CAAAATATAAGTGCTAACAG**TCGTCTGCAAGTTTAG | Overlap |
| Syn_PartB_R | **CCCAGTTGGGG**TCGACTCTAG | Overlap |
| Vec_F | CCCCAACTGGGGTAACCTTTG | Overlap, vector linearization |
| Vec_R | TCGGCTTAATAGCTCACGCTATGC | Overlap, vector linearization |
| INS_vec_PartA_F | CTTTAAAATGCCCCTTAAAATTCAAAATAAAGGC | Plasmid insertion check |
| INS_PartAB_R | TACTAGATGTTGGCAAACGATATGAGC | Plasmid insertion check |
| **Assembly of DNA editing templates for recombineering (insertion of GAGG MTase)** | | |
| Dcm_UP_FW | **AAATAAGGTTAACAT**TCGGTAAGCGCTTCATCCG | Overlap |
| Dcm_UP_RV | **TTTAAATT**CCATGGCCACTCGCAGCAAAAATATG | Overlap |
| Dcm_DOWN_FW | **GAATTTGATATTTAA**AGATTTCACCGGCCATCTG | Overlap |
| Dcm_DOWN_RV | **CGGGGATCCTGTCC**AGGATGCGGATCG | Overlap |
| MTase_FW | **GCGAGTGGCC**ATGGAATTTAAAATATTAGATTTG | Overlap |
| MTase_RV | **CCGGTGAAATCT**TTAAATATCAAATTCTAAGTTAG | Overlap |
| pRRS_Vector_FW | **CATCCTGGACAG**GATCCCCGGGGAAGATC | Overlap |
| pRRS_Vector_RV | **TGAAGCGCTTACCGA**ATGTTAACCTTATTTCCTGC | Overlap |
| Plas_ins_up_F | ATTAATGTGAGTTAGCTCACTCATTAGG | Plasmid insertion check |

| | | |
|---|---|---|
| Plas_ins_down_R | AATACCGCACAGATGCGTAAG | Plasmid insertion check |
| pRRS_Dcm_51-13_ET_FW | TCGGTAAGCGCTTCATCCGTCAGC | Produces linear editing template |
| pRRS_Dcm_51-13_ET_RV | TGTCCAGGATGCGGATCGGCTG | Produces linear editing template |
| **Knockouts using pIMAY-Z** | | |
| IM3-R | AATACCTGTGACGGAAGATCACTTCG | Insertion check. Monk and Stinear (1) |
| IM4-F | TACATGTCAAGAATAAACTGCCAAAGC | Insertion check. Monk and Stinear (1) |
| T3_FW | GCATGTAAAAAACAAGCGGGCTTTGC | TI target removal |
| T3_RV | CAATACAATGTAGGCTGCTCTACAC | TI target removal |
| SecA-1-R | **AACACCTCATT**ATTAGCCATTTTGATCACCTCG | Overlap |
| SecA-1-F | **ACAAAAGCTG**TACTACAACCTTTATGTATGGTACGAAATTACGCA | Overlap |
| SecA-2-F | **CAAAATGGCTAAT**AATGAGGTGTTCATAGATGACAATATA | Overlap |
| SecA-2-R | **GGGCGAATTGT**ATAATTTGGCACGATTCTCATATG | Overlap |
| SecA-VEC-F | **CCAAATTATACAA**TTCGCCCTATAGTGAGTCGT | Overlap, vector linearization |
| SecA-VEC-R | **GGTTGTA**GTACAGCTTTTGTTCCCTTTAGTG | Overlap, vector linearization |
| SecA-KOcheck-F | TTATCACCAGTATTGAGTCAACAAGG | Genome deletion check |
| SecA-KOcheck-R | GTACAACAATACCGACTTTACTTGATC | Genome deletion check |
| SraP_T1_FW | GAGGCTAAAGTATAATAATTGGTTGGAATAATGTATG | TI target removal |
| SraP_T1_RV | TTTTTTTAAGGATAGACAACCAATTTATAGAAAAG | TI target removal |
| SraP-1-F | **CAAAAGCTGG**TTTTTTGATTAATTTTTATTTTTC | Overlap |
| SraP-1-R | **GT**AATACTCTCCTTTATTCTCACACTTAC | Overlap |
| SraP-2-F | **GGAGAGTATT**ACTACTTCATGTGGGAGATAG | Overlap |
| Srap-2-R | **GCGAATTGG**TTATAATTCTTTCCTTATAATGC | Overlap |
| SraP-VEC-F | **GAATTATAAC**CAATTCGCCCTATAGTGAGTCG | Overlap, vector linearization |
| SraP-VEC-R | **CAAAAAAC**CAGCTTTTGTTCCCTTTAGTGAG | Overlap, vector linearization |
| SraP-KOcheck-F | GTTATGGCTACAACAATGAGGGAGAG | Genome deletion check |
| SraP-KOcheck-R | GTGTTGGTTATTAAATAATGATTTGATTAAACTTGTTAATAC | Genome deletion check |
| CapA_DW_FW | **CGAAATTAGAATTTG**CATGGCTAAAAATAAAAAAGAAGTAAG | Overlap |
| CapA_DW_RV | **GCGAATTG**CATCTGTCACAGTGTTTACTGGT | Overlap |
| CapA_UP_FW | **GAACAAAAGCTG**ATAAATGTAGAACCTTAAAGGAGTGAC | Overlap |
| CapA_UP_RV | **GCCATG**CAAATTCTAATTTCGATATTAATTTTATATTTTTAACATGG | Overlap |

| | | |
|---|---|---|
| CapA_Vec_FW | **GACAGATG**CAATTCGCCCTATAGTGAGTCGTAT | Overlap, vector linearization |
| CapA_Vec_RV | **TTCTACATTTAT**CAGCTTTTGTTCCCTTTAGTGAGGG | Overlap, vector linearization |
| CapA_KO_Check_FW | GCTAATTCATCTAATGGGTGAGCTGTAATG | Genome insertion check |
| CapA_KO_Check_RV | CTTCCTTACTTTTAGGACCGTCATCTAC | Genome insertion check |
| CapA_T1_FW | CTCTGTGTAGAAATCTCTTGGCTTTTTTATATTTATG | TI target removal |
| CapA_T1_RV | ATATCTTGGCTTAAAACATTAATATTCAATATTTGGC | TI target removal |
| CapI_DW_FW* | **AAGGATAAAAAA**ATATAGTGGTGAGAAAATGAGTAAATCCAAAGATTT**C**ATATTTAG | Overlap, TI target removal |
| CapI_DW_RV | **GGCGAATTG**CTGAAAATATTGAAACATCATAGAAGTACAC | Overlap |
| CapI_UP_FW | **AACAAAAGCTG**GATAATGAAAAAAACTTCCTGTCACTAATGAATGC | Overlap |
| CapI_UP_RV | **CTCACCACTAT**ATTTTTTTATCCTTTCTTTACACTATTCATCAAAG | Overlap |
| CapI_Vec_FW | **CAATATTTTCAG**CAATTCGCCCTATAGTGAGTCGTATTACG | Overlap |
| CapI_Vec_RV | **TTTTCATTATC**CAGCTTTTGTTCCCTTTAGTGAGG | Overlap |
| CapI_KO_Check_FW | CACGGAACATAGTAGACGCTATTTATTAGATG | Genome insertion check |
| CapI_KO_Check_RV | CATCATTTACTTCGTAGCTTAAAAGTACTGTTAC | Genome insertion check |
| Epsn_INS_FW | GTGATTGAATGTGTAAAACTTTTTTATCTGTTAAAAG | Overlap |
| Epsn_INS_RV | CGTTGATGACCGTGATAAGAAAGAC | Overlap |
| Epsn_DW_FW | **AATCTTAACTAA**ATGAAAGCACTTTGTCGATGATATAAATC | Overlap |
| Epsn_DW_RV | **GGGCGAATTG**TTATCTAAACTAGAAAAGAATGATAAAGGGTTCTTCTTAATGG | Overlap |
| Epsn_MID_FW | **GCAAGTAGTTTATTTATTACATATTTAC**ATGATAGTTACATTAGTTTTAACGTAAATTCTAAC | Overlap |
| Epsn_MID_RV | **AAGTGCTTTCAT**TTAGTTAAGATTTAAGTTTTCTTTTAATTTCTCATTAACT | Overlap |
| Epsn_UP_FW | **CAAAAGCTG**CATCTAATGGGTGAGCTGTAATGTATAATAATTTTGC | Overlap |
| Epsn_UP_RV | **GTAACTATCATGTAAATATGTAATAAATAAACTACTTGC**GTAATGAATTAGTAGC | Overlap |
| Epsn_vec_FW | **GTTTAGATAA**CAATTCGCCCTATAGTGAGTCGT | Overlap, vector linearization |
| Epsn_vec_RV | **CCATTAG**ATGCAGCTTTTGTTCCCTTTAGTGAGGG | Overlap, vector linearization |

**Supplementary Table S2) Plasmids used in this study**

| Plasmid | Addgene number | Reference |
|---|---|---|
| pRRS | | Johnston, Cotton (2) |
| pEPSA5 | | Johnston, Cotton (2) |
| pTarget | #62226, modified by Johnston et al. for *Dcm* target | Jiang, Chen (3), Johnston, Cotton (2) |
| pCas | #42876 | Jiang, Chen (3) |
| pIMAY-Z | | Monk and Stinear (1) |

**Supplementary Table S3) Bacteria strains used in this study**

| Strain/organism | Description/Accession number | Reference |
|---|---|---|
| *S. haemolyticus* 8-9 | SAMEA1035041 | Cavanagh, Klingenberg (4) |
| *S. haemolyticus* 25-63 | SAMEA1035076 | Cavanagh, Klingenberg (4) |
| *S. haemolyticus* 51-13 | SAMEA1035044 | Cavanagh, Klingenberg (4) |
| *S. haemolyticus* 51-29 | SAMEA1035047 | Cavanagh, Klingenberg (4) |
| *S. haemolyticus* 51-41 | SAMEA1035075 | Cavanagh, Klingenberg (4) |
| *S. haemolyticus* 53-27 | SAMEA1035140 | Cavanagh, Klingenberg (4) |
| *S. haemolyticus* 53-38 | SAMEA1035061 | Cavanagh, Klingenberg (4) |
| *S. haemolyticus* 53-49 | SAMEA1035106 | Cavanagh, Klingenberg (4) |
| *S. haemolyticus* 57-1 | SAMEA5568753 | Cavanagh, Klingenberg (4) |
| *S. haemolyticus* 51-22 | SAMEA5568761 | Cavanagh, Klingenberg (4) |
| *E.coli* JMC1 | Minicircle producing *E. coli* | Johnston, Cotton (2) |
| *E.coli* JMC4 | Minicircle producing *E. coli* mimicking *S. haemolyticus* MTase. | This study |
