## Supplementary materials for "Genetic engineering of *Staphylococcus haemolyticus*: Overcoming Restriction-Modification Barriers and Targeting Virulence Genes"

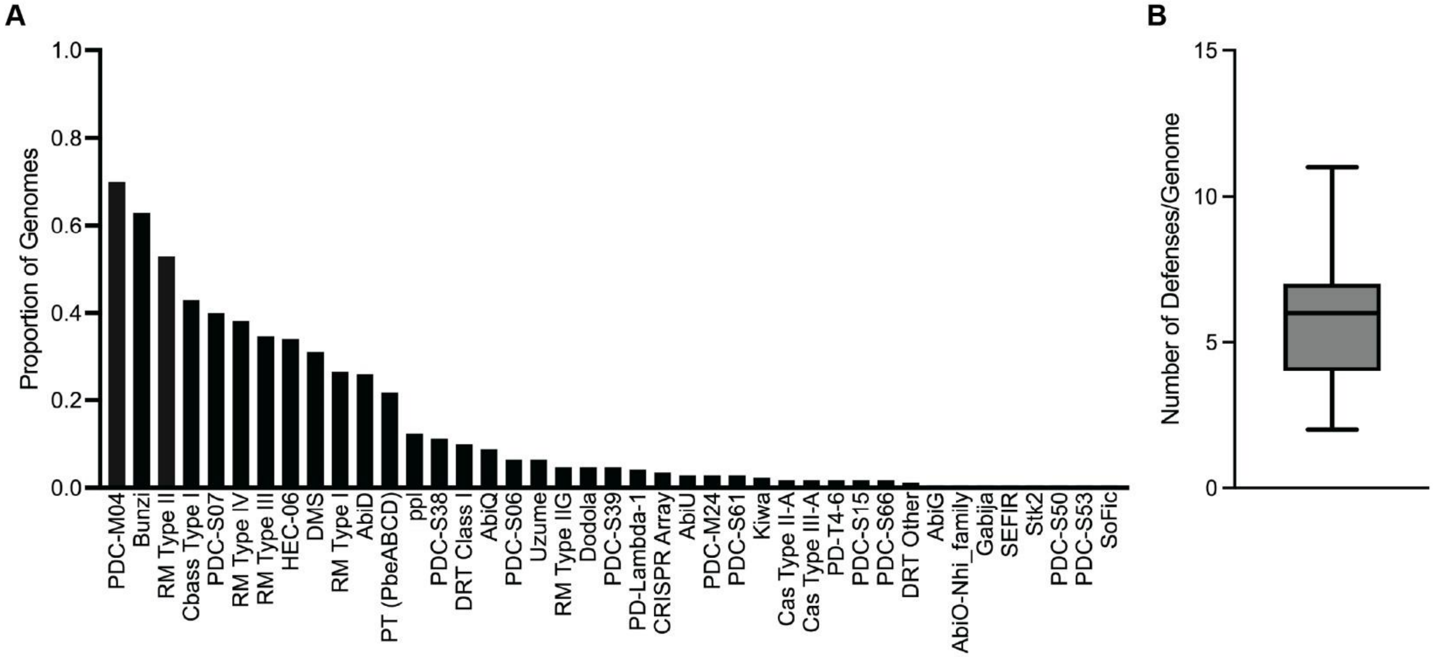


Figure S1) Number of genomes which contain each of the defense systems detected (**A**), and number of defense systems per genome (**B**). PDC, Phage defence candidate; RM, restriction modification; Cbass, cyclic-oligonucleotide-based antiphage signalling system; Hec, Hma-embedded candidate; DMS, DNA-modification system; Abi, abortive infection; PT, Phosphorotioation system; ppl, polymerase/histidinol phosphatase-like; DRT, defense-associated reverse transcriptase; PD, phage defence; CRISPR, clustered regularly interspaced short palindromic repeats; Cas, CRISPR-associated protein, Stk, Serine/threonine kinase; SoFic, standalone protein with Fic domain.

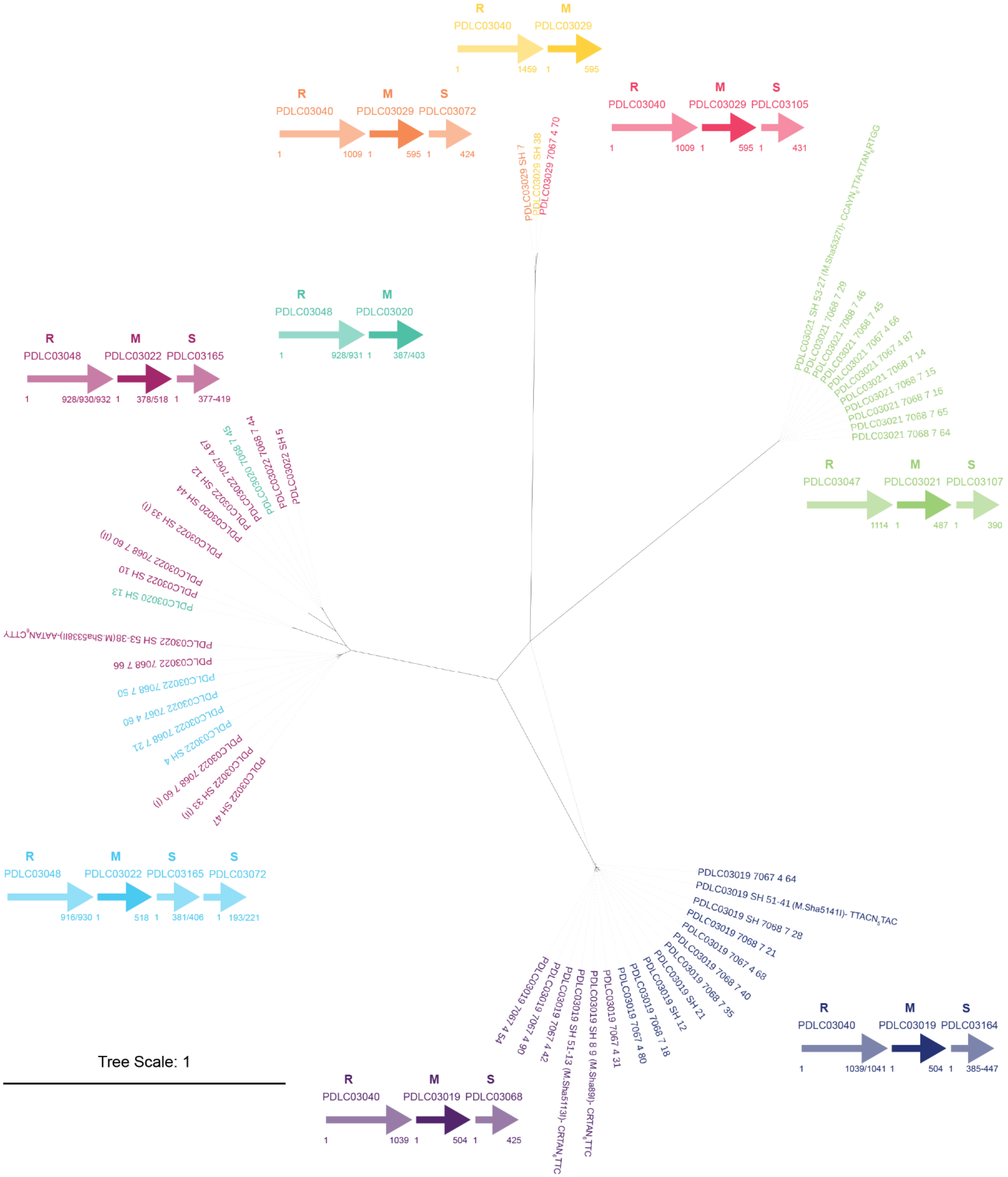


Figure S2A Type I MTases detected in *S. haemolyticus* genomes

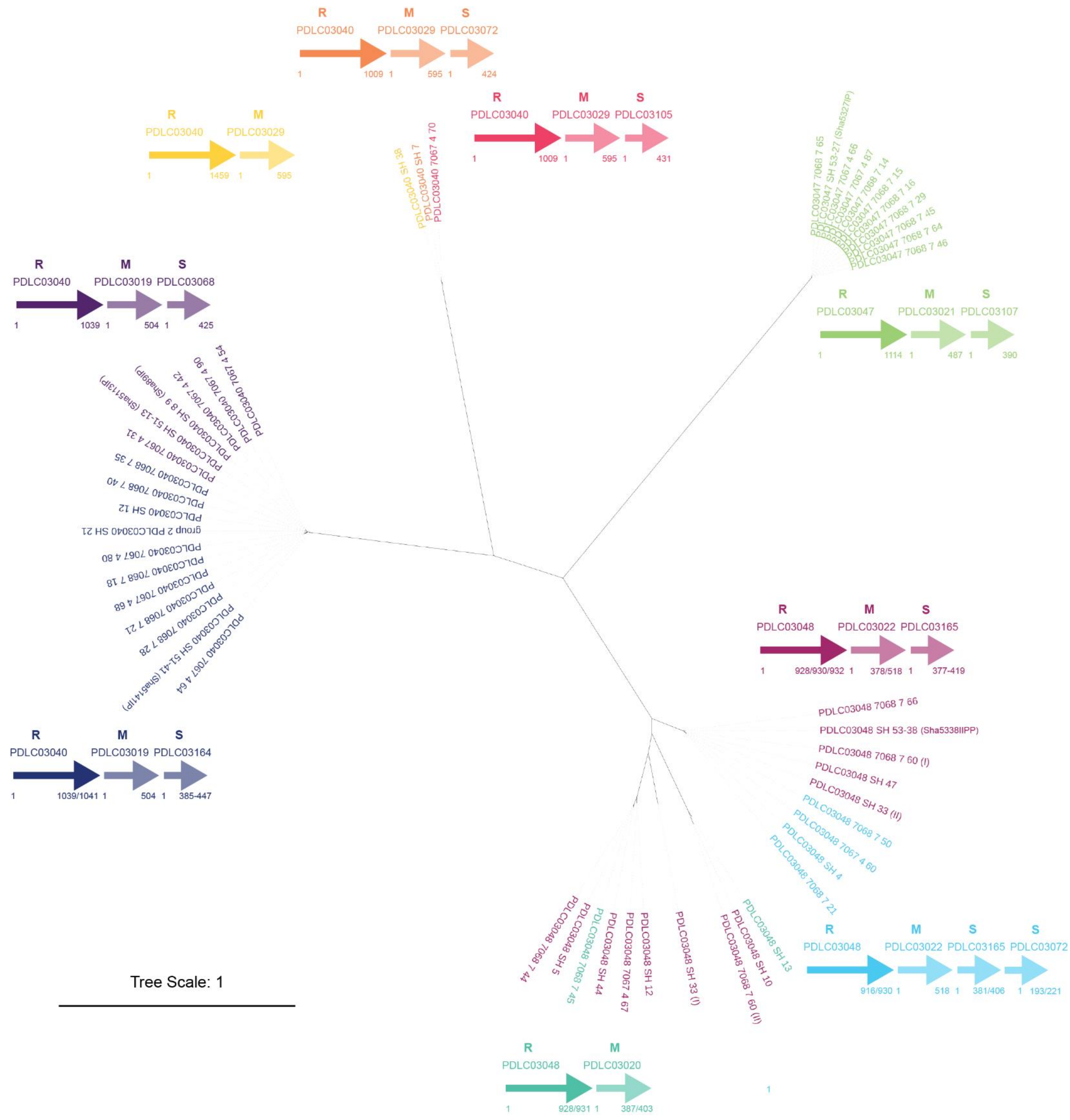

Figure S2B Type I REases detected in *S. haemolyticus* genomes

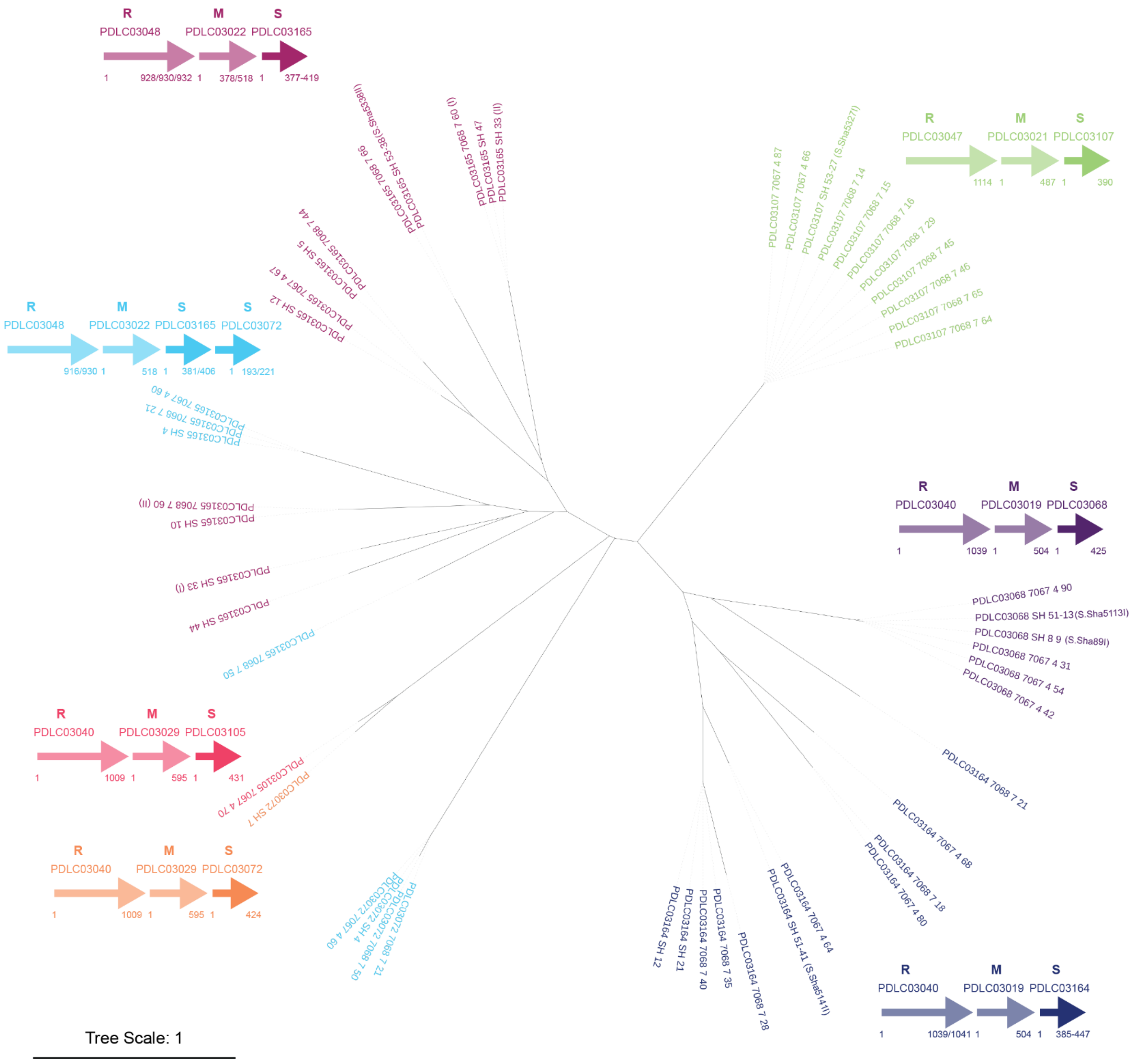


Figure S2C Type I Specificity subunits detected in *S. haemolyticus* genomes

Figure S2A-C) Maximum-likelihood dendrograms based on MUSCLE-aligned amino acid sequences of Type I system proteins identified by PADLOC. Branches represent proteins with highly similar sequences. Leaf labels show PADLOC accession number and the strain identifier for the genome in which the protein was detected. PADLOC and REBASE data was matched (Supplementary Table X) and, when available, the REBASE-assigned methylation motif is also included. Colours indicate proteins which occur together as part of the same system. The arrangement of the genes in each system is also given; bold arrows correspond to the elements present on the tree. R, restriction enzyme (REase), M, methyltransferase (MTase), S, specificity subunit.

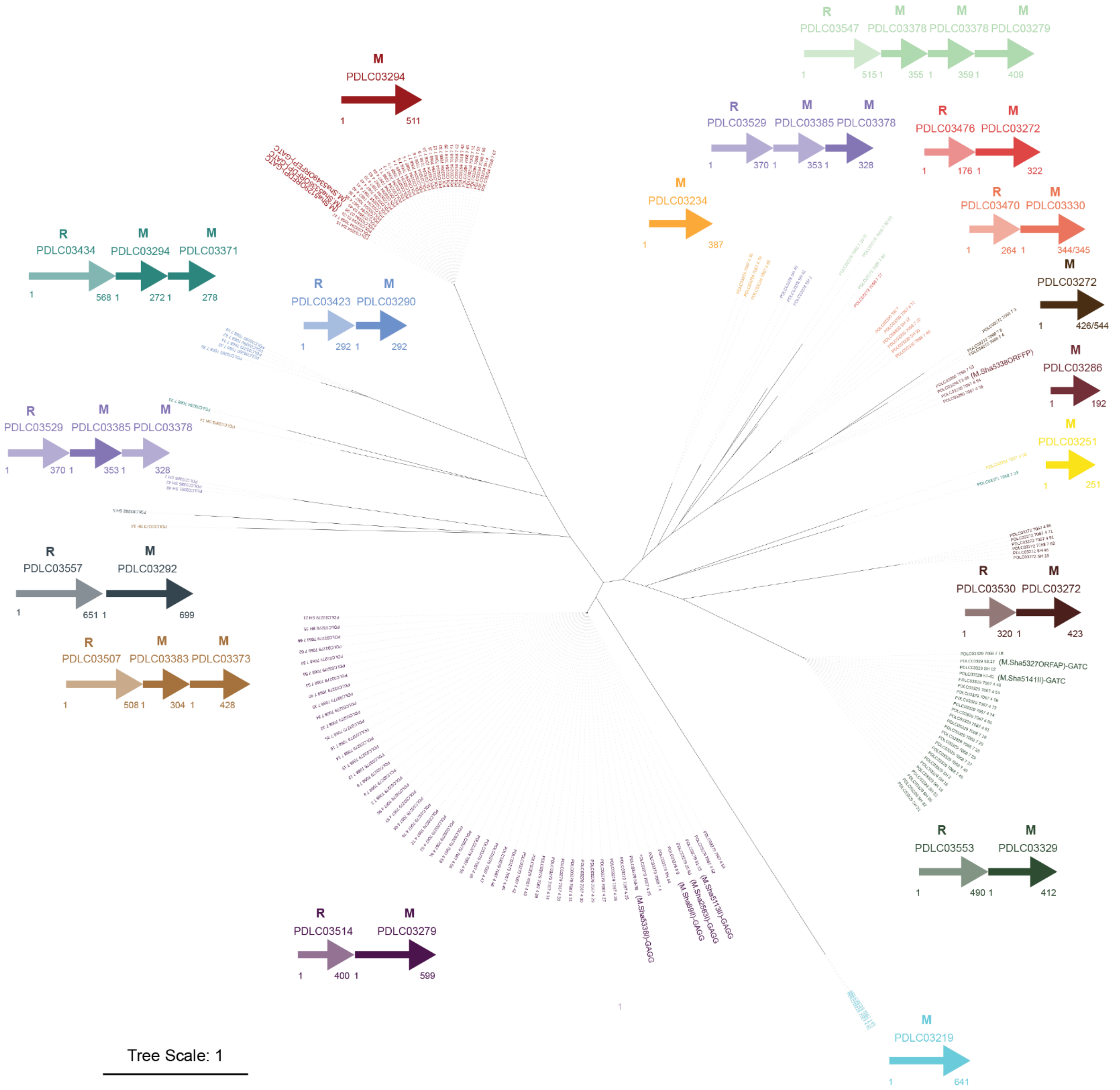


Figure S3A Type II MTases detected in *S. haemolyticus* genomes

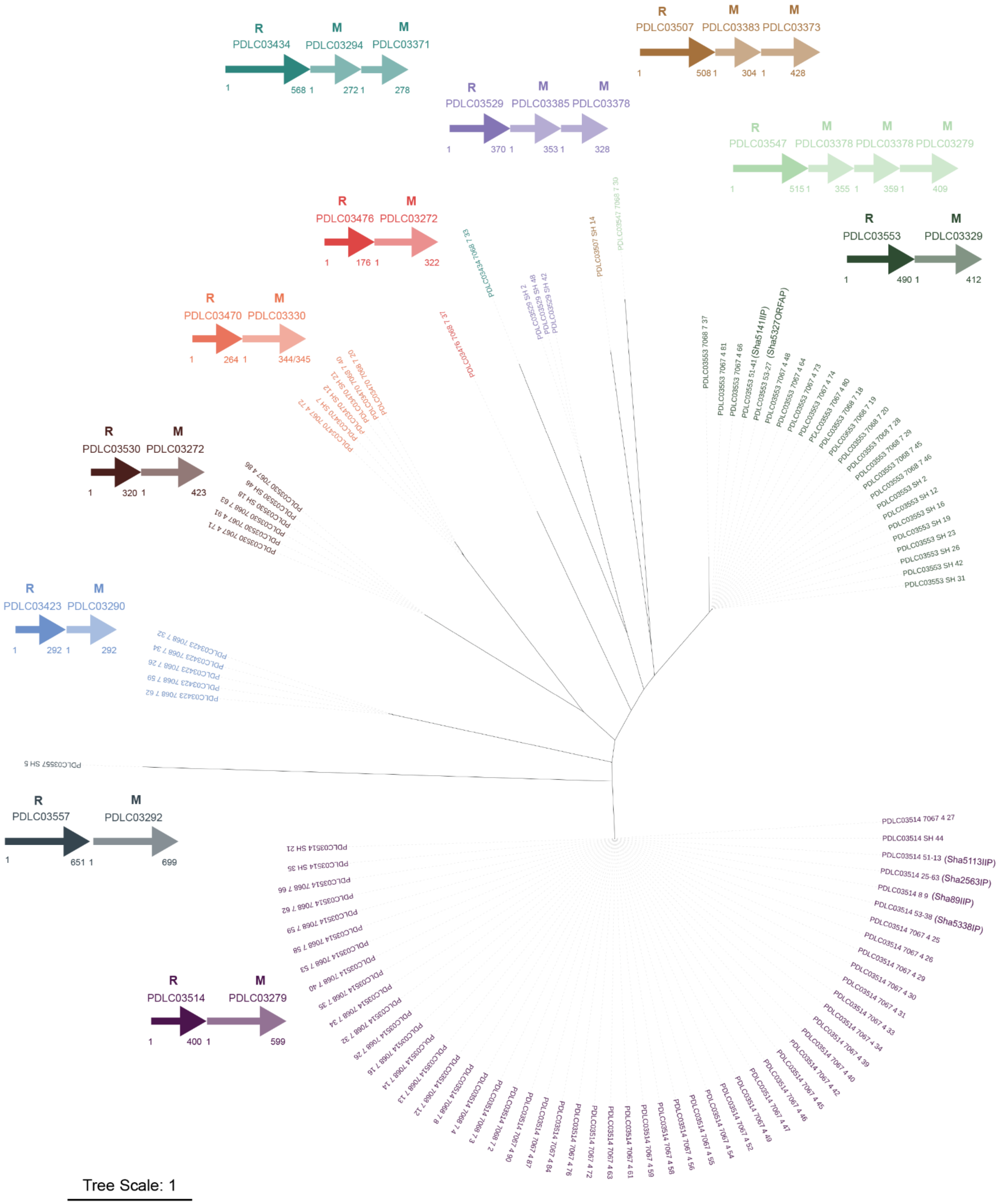


Figure S3B Type II REases detected in *S. haemolyticus* genomes

Figure S3A-B) Maximum-likelihood dendrograms based on MUSCLE-aligned amino acid sequences of Type II system proteins. Branches represent proteins with highly similar sequences. Leaf labels show PADLOC accession number and the strain identifier for the genome in which the protein was detected. PADLOC and REBASE data was matched (Supplementary Table X), and when available, the REBASE-assigned methylation motif is also included. Colours indicate proteins which occur together as part of the same system. The arrangement of the genes in each system is also given; bold arrows correspond to the elements present on the tree. R, restriction enzyme (Rease), M, Methyltransferase (MTase).

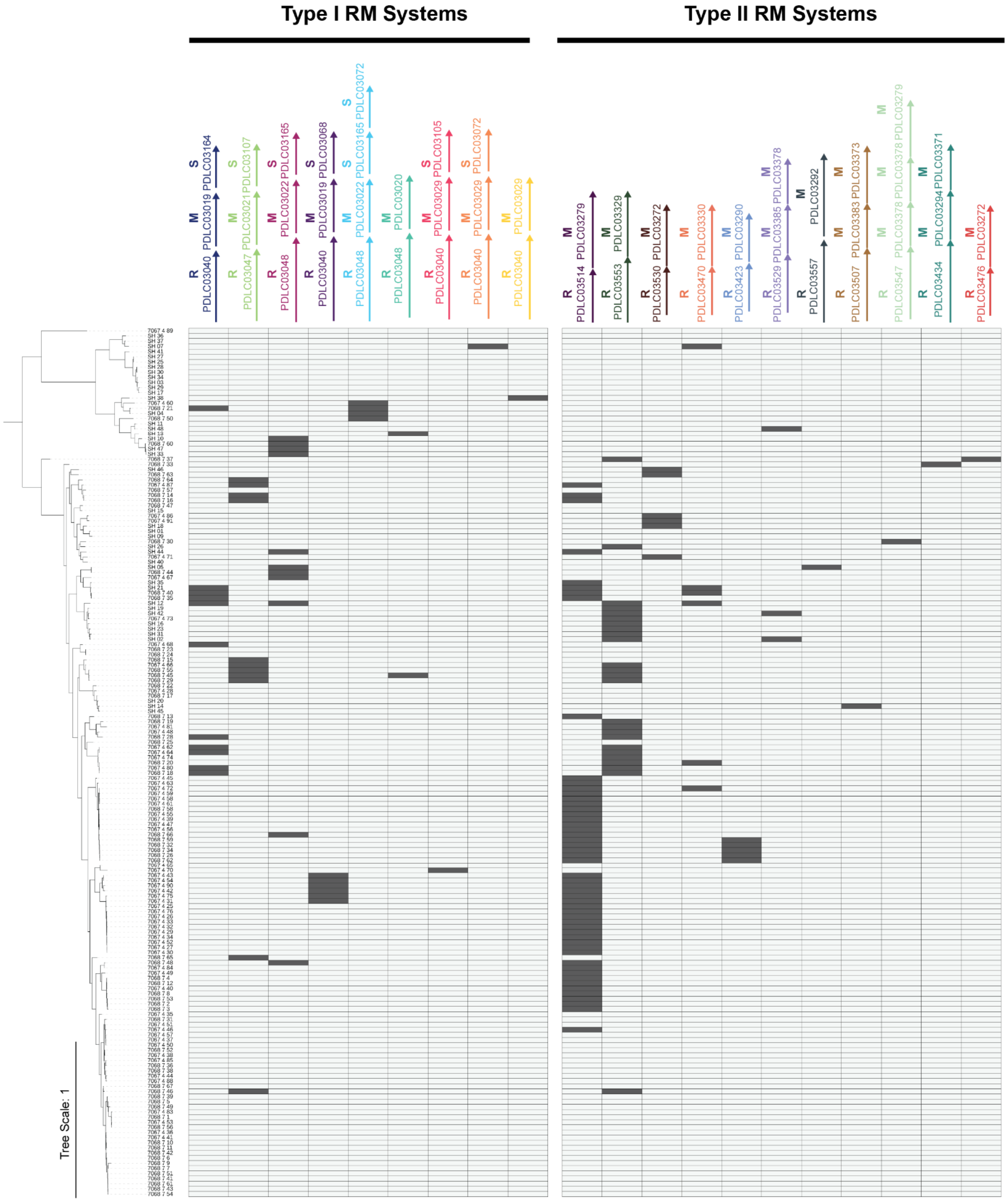


Figure S4) A reference-free phylogenetic tree constructed from whole genome sequences showing the distribution of Type I and Type II systems across *S. haemolyticus* isolates
