## Supplementary methods for "Genetic engineering of *Staphylococcus haemolyticus*: Overcoming Restriction-Modification Barriers and Targeting Virulence Genes"

**Genomic DNA isolation for high-purity, high molecular weight DNA**

High molecular weight (HMW) genomic DNA was isolated using a modified protocol based on the MasterPure™ Gram Positive DNA Purification Kit (Lucigen). Bacteria were grown overnight in 10 ml of TSB, at 37°C, with shaking at 250 rpm. The overnight cultures were pelleted by centrifugation at 5000 rpm for 10 minutes. The supernatant was removed and the tubes were inverted onto tissue paper to draw out residual media. Resuspend the cells in at least 10 ml TES buffer TES buffer (20 mM Tris-HCl, pH 8.0, and 50 mM EDTA) and normalize the sample to OD600 of 1. 10 ml of the normalized cell suspension were transferred to a new centrifuge tube and pelleted at 5000rpm for 10min. The supernatant was removed and the cells resuspended in 300 µL of TE buffer (from the kit). The suspension was transferred to Lysing Matrix B tubes, and vortexed briefly before 1 µL of lysozyme was added. Samples were vortexed briefly, pulse-spun, and incubated at 37°C for 20 minutes. Following this, 300 µL of GP lysis solution was added, and the samples were bead-beaten at 6.0 m/s for 20 seconds using an MP FastPrep®-24 instrument. For tougher strains, additional bead-beat cycles may be performed with 5-minute cooling intervals on ice.

Proteinase K (2 µL) was added to each sample, followed by incubation at 65°C for 15 minutes. Samples were cooled to 37°C for 5 minutes, placed on ice for 3–5 minutes. 350 µL of MPC Protein Precipitation Reagent was added, then mixed by pulse vortexing for 3-5 seconds before placing back on ice. Debris was pelleted by centrifugation at 10 000 g for 10 minutes at 4°C. The supernatant was carefully transferred to a clean1.5 ml Eppendorf tube, centrifuged again for 2 minutes at 10 000 *g*, and transferred to a new tube to minimize debris carryover. RNase A (2 µL) was added, and samples were incubated at 37°C for 30 minutes.

DNA was precipitated by mixing the samples with 700 µL of DNA-grade isopropanol, inverting 30–40 times, and centrifuging at 10,000 rpm for 10 minutes at 4°C. The resulting DNA pellet was washed twice with 1 ml of 70% ethanol (the pellet was carefully dislodged from the side of the tube by gentle pipetting or flicking the tube, then the tube was inverted several times to ensure removal of the salts). The ethanol was removed, and the tubes were briefly centrifuged before the remaining ethanol was removed by pipette. The pellets were air-dried, then resuspended in 50–200 µL of pre-warmed TE buffer or water at 37°C. If the pellets were tricky to resuspend, the tubes were incubated at 37°C in a dry bath until they became translucent and flicked gently.

DNA quality was assessed using a Nanodrop spectrophotometer (A260/280 and A260/230 ratios), a Qubit fluorometer for dsDNA quantification, and agarose gel electrophoresis to confirm HMW integrity. Only DNA with a yield of 10–15 µg and minimal fragmentation was used for PacBio sequencing.

**Syngenic DNA**

Isolate 51-13 has the methylation motifs GAGG/CCTC (Type II) and CRTANNNNNNNTTC (Type I). We used plasmid pEPSA5, which had previously been modified to have minicircle-forming properties(1). pEPSA5 did not contain any CRTANNNNNNNTTC motifs, but did have 25 GAGG/CCTC motifs, 17 of which were located on the *E. coli* replicon and were not edited because they would be removed during minicircle formation. The remaining 8 motifs were located in the *S. aureus* replicon. We edited the plasmid sequence using DNAstar. When the motifs fell within coding sequences, we aimed to create silent mutations that would not change the amino acid codon in that position. Two motifs were located within the promoter region of the chloramphenicol resistance gene; these we left unaltered. These edited sequences were sent to Synbio Technologies (Monmouth Junction, USA) to be synthesised. The synthesised fragments and plasmid backbone were used as templates for PCR in which primers contained overlaps needed for assembling the fragments into circular plasmids. After cleaning the PCR products and denaturing template DNA using DpnI (New England Biolabs, USA), we used NEBuilder Hi-Fi Assembly Master Mix (New England BioLabs, USA) to assemble the plasmids. Assembled plasmids were transformed into NEB 5-alpha Competent *E. coli*. Colony PCR (see supplementary table 1 for primers) was done the following day to select correctly assembled plasmids, which were isolated using the NucleoSpin Plasmid kit (Macherey Nagel, country). The minicircles were linearised with restriction enzyme HindIII (New England BioLabs, USA) to check for residual full plasmids.

Table S8. Composition of the plasmids and minicircles constructed for transformation of S. haemolyticus isolate 51-13. The suffix "Syn" denotes a syngenic construct in which GAGG sites have been edited out, while the "MC" suffix denotes a minicircle, which is induced to lose the *E. coli* replicon borne on pEPSA5.

| Construct | Size (bp) | Number of GAGG sites |
|---|---|---|
| pEPSA5 | 8361 | 25 |
| pEPSA5MiniMC | 2668 | 8 |
| pEPSA5MiniSynMC | 2668 | 2 |

### Simplified *E. coli* transformation

Isolated plasmids were electroporated into JMC1, an *E. coli* strain previously modified to produce minicircles (1, 2). A simplified electroporation protocol was used (3): JMC1 was grown on LB agar at 37°C overnight. A small amount of culture (about 2 µL) was picked off the plate with a sterile 1 µL inoculation loop and placed into 700 µL of sterile Milli-Q water in a 1.5 ml Eppendorf tube, then vortexed to resuspend. The cells were pelleted at 8000 *g* for 3 minutes; then the supernatant was removed. The cells were resuspended in 700 µL of sterile Milli-Q water and pelleted again at 8000 *g* for 3 minutes. The supernatant was removed, and the cells were resuspended in 40 µL of sterile Milli-Q water. The plasmids or minicircles were added (approximately 100 ng) to the cells and mixed gently before being transferred to a 0.1 cm electroporation cuvette (Biorad). The cells were electroporated at 1.8 kV, 200 Ω and 25 µF (Biorad Gene Pulser Xcell Microbial System). After electroporation, 1 ml of pre-warmed LB broth was added to the cells before transfer to a fresh Eppendorf tube. The cells were incubated at 37°C with gentle shaking for 1 hour before being plated out on LB agar containing 50 µg/ml kanamycin and incubated overnight. Colony PCR was done to confirm the presence of the plasmid/minicircle.

### Minicircle induction

A single transformant colony was selected and used to inoculate 5 ml Terrific Broth (TB) containing 50 µg/ml kanamycin and incubated with shaking at 250 rpm at 30°C for 4-6 hours. This pre-culture was diluted 1:200 in fresh TB broth + 50 µg/ml kanamycin in an Erlenmeyer flask (5 × the volume of the broth), and incubated with shaking at 250 rpm at 30°C overnight (but not more than 16 hours).

The pH and OD600 of the overnight culture were measured. If the OD600 is > 8 with pH lower than 6,5, check the ventilation of the incubator and start over. If the pH was above 6.5, and the OD600 was between

4 and 6, the overnight culture was combined with the same volume of induction media, and if the OD600 was between 6 and 8, the volume of induction media added was double that of the overnight culture. Induction media was made up of 400 ml LB broth, 16 ml 1N NaOH and 400 µl 20% Arabinose, filter sterilised. A 1 ml aliquot (time 0, pre-induction) was removed for plasmid isolation. The induction mix was then incubated at 30°C with shaking at 250 rpm for 3 hours, and then at 37°C for 1 hour. Another 1 ml aliquot was removed for plasmid isolation and run on a gel alongside the time 0 pre-induction plasmid. If induction was not complete, the flask was returned to the 30°C and incubated with shaking for another hour. When induction was completed, the bacteria were pelleted by centrifuging at 5000 *g* for 10 minutes. The supernatant was removed, and the pellets either underwent minicircle isolation immediately using the NucleoSpin Plasmid kit (maxi prep), or they were frozen at -20°C until minicircle isolation could be done.

The full-sized plasmids and minicircles were used to optimise the transformation of *S. haemolyticus* 51-13 by tweaking the conditions used to produce competent cells, electroporation conditions, and recovery conditions.

***S. haemolyticus* competent cell preparation and electroporation**

For preparing competent cells, a saturated overnight culture of *S. haemolyticus* was diluted to an OD600 of 0.25 with TSB and incubated in an Erlenmeyer flask (5x the volume of broth used) at 37°C, with shaking at 250 rpm. When the OD600 reached 0.8-1.0, the flask was placed in an ice slurry for 15 minutes. The cells were harvested by centrifuging at 5000 *g* for 10 minutes at 4°C (all centrifuging was done at 4°C). The cells were washed in an equal volume of ice-cold $ddH_2O$, then pelleted again at 5000 *g* for 10 minutes. The wash step was repeated, then after pelleting, the cells were resuspended in 1/10 volume of 10% glycerol (ice cold). The cells were pelleted again, then resuspended in 1/25 volume 10% glycerol. The cells were concentrated in this stepwise fashion using 1/50 and then 1/100 volumes. Finally, the cells were resuspended in 1/165 volume and aliquoted into suitable volumes and frozen at -80°C until use.

For electroporation, the aliquots were thawed on ice (5 minutes) and then on the bench (5 minutes). The cells were pelleted by centrifuging at 5000 *g* for 1 minute, followed by removal of the supernatant. The cells were resuspended in the same volume of 10% glycerol + 0.5 M sucrose and split into 50 µL aliquots in 1.5 ml microcentrifuge tubes. 1-5 µg of the plasmid/minicircle was added and mixed gently by pipetting up and down 2-3 times, before incubation on the bench for 10 minutes. The plasmid and cell mix was transferred to a 1 mm electroporation cuvette (Biorad) and pulsed at 2,5 V, 100 Ω, 25 µF (Biorad Gene Pulser Xcell Microbial System). 950 µL of recovery buffer (TSB + 0.5 M sucrose) was added, and the cells were incubated at 37°C with gentle rotation (250-300 rpm) for 1 hour before plating the bacteria out on selective media and incubating for 24-48 hours (depending on the plasmid used).

**Constructing JMC4**

Assessing our *S. haemolyticus* collection, we found that 55 isolates had a methylation system identical to the one in 51-13 that produced the GAGG/CCTC motif. This, in combination with the tendency of the GAGG/CCTC motif to fall within regulatory regions (making it difficult to edit out), led to our decision to construct an *E. coli* strain to replicate the GAGG/CCTC pattern in passaged plasmids. We used the method described by Jiang, Chen (4) to knock the *S. haemolyticus* MTase gene into JMC1 *E. coli*, while simultaneously knocking out the native *dcm* gene (a MTase methylating the CCWGG motif).

We constructed the editing template by using primer set in Table S1 to amplify the MTase gene in 51-13, and to amplify the upstream and downstream regions of the *dcm* gene from JMC1. The three sections were assembled into plasmid pRRS using the NEBuilder Hi-Fi Assembly Master Mix, with the MTase gene positioned between the two *E. coli* flanking regions. This construct (editing template) was linearized by PCR.

JMC1 was transformed with the pCas plasmid, which contained the λ-Red genes in addition to the Cas9 gene. JMC1+pCas electroporated with 100 ng of plasmid pTarget (obtained from Johnston et al, already carrying the sgRNA targeting dcm) and 400 ng of the editing template. After recovery at 30°C, the bacteria were plated out on LB agar containing kanamycin and spectinomycin and incubated at 30°C overnight. Confirmation of the insertion was determined by PCR and whole genome sequencing (sequencing and methylation analysis done in-house by the Johnston lab). DNA from the transformants was also digested with restriction enzyme MnII to ascertain whether the GAGG/CCTC motif was successfully methylated (see supplementary results 1). The resulting strain was named JMC4, and it is able to replicate the GAGG/CCTC methylation pattern and produce minicircles.

**Hybrid approach**

It is possible to combine the mimicry and SynGenic strategies. When plasmids were passaged through JMC4, it was not necessary to edit out the GAGG motifs. However, some plasmids contained motifs of type I systems. Since the motifs did not occur many times on any of the plasmids, we employed site-directed mutagenesis instead of DNA synthesis to edit out these motifs. Plasmids were amplified using primers designed to replace individual nucleotides in order to disrupt the type I restriction target sequence. Amplification was done using Q5 2X Mastermix, and the product was used as a template for a KLD reaction (New England BioLabs), resulting in a re-circularised plasmid without the type I system motifs. The plasmids were then passaged through JMC4 before being transformed into the target strains.

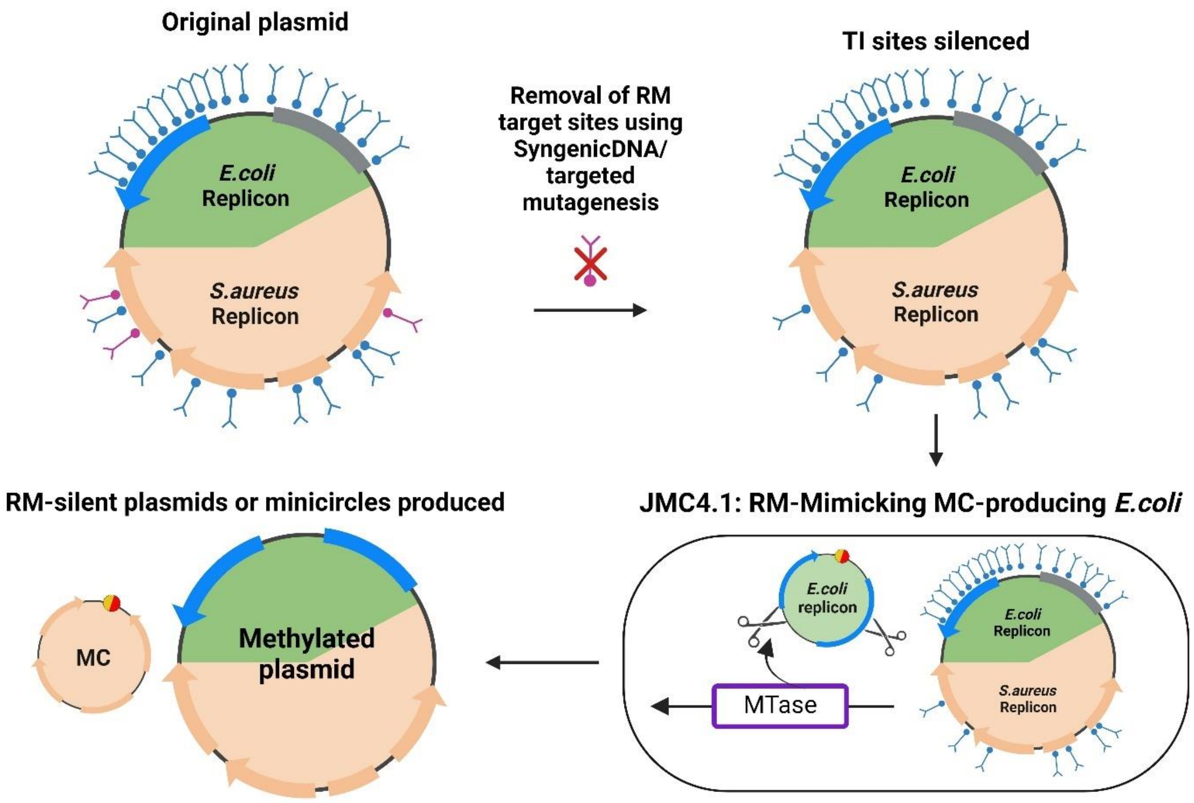


Figure S5: Combining mimicry and Syngenic strategies to create RM-silent plasmids for transformation into *S. haemolyticus*

**Allelic exchange**

Plasmid pIMAY-Z carried chloramphenicol as a resistance marker and the Lac-Z operon which allows for phenotypic screening of plasmid integration and excision. It works by allelic exchange, which uses homologous recombination to incorporate a template that excludes the gene to be deleted. We targeted several genes in strain 53-38 for deletion: *sraP*, *secA2*, *capA* and *capA*. We also replaced the capsule operon with the genes *EpSN* and *tagU*.

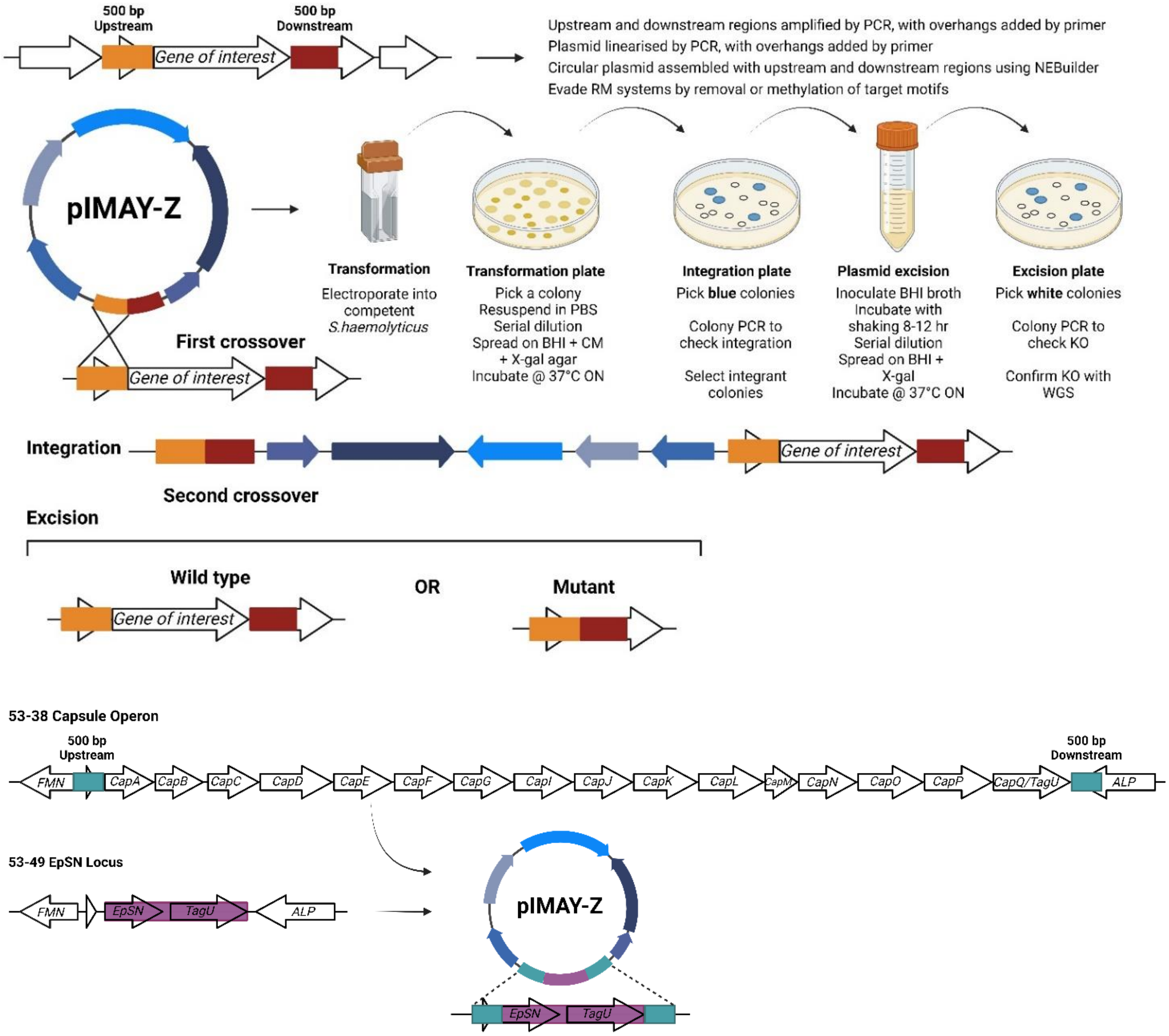


Figure S6: Summary of the allelic exchange process and construction of the insert for simultaneous deletion of capsule operon and insertion of the *EpSN* and *TagU* genes.

We followed the method set out in Monk and Stinear (5), except for the following changes: We used the NEBuilder Hi-Fi Assembly Master Mix to assemble the plasmid, instead of SLiCE, and adapted the methods for producing competent cells and transformation for *S. haemolyticus* (described above). Type I restriction sites on the plasmids were removed using site-directed mutagenesis, and plasmids were passaged through JMC4.1 to mimic GAGG/CCTC methylation.

For each of the genes of interest, a version of the pIMAY-Z plasmid containing a template for allelic exchange was constructed using NEBuilder® HiFi DNA Assembly. In each case, the recombination template was made by selecting approximately 500 bp upstream and downstream of the gene of interest, amplifying them by PCR, using Q5® High-Fidelity 2X Master Mix (New England Biolabs) and genomic DNA from 53-38 as a template. The plasmid was linearized by PCR. The primers belonging to fragments which would be adjacent to each other when the vector was assembled contained matching overlapping sequences at their 5' ends.

In non-capsulated *S. haemolyticus* isolates, a different gene *EpSN*, as well as *tagU* are present in the locus where the capsule operon is found in capsulated strains. For this experiment, instead of deleting the capsule operon, the operon was replaced by the *EpSN* gene and the *TagU* cell envelope-associated transcriptional attenuator gene which was located immediately downstream. This region was amplified from 53-49, a non-capsulated isolate. The regions upstream and downstream of the capsule operon in isolate 53-38 were also amplified, and all three fragments were assembled into the pIMAY-Z plasmid. The plasmid was then treated in the same manner as the other allelic exchange vectors described above.

We did not obtain any direct integrants, so followed the slow integration protocol following transformation (5, 6), as illustrated in figure S6. Briefly, a colony which appeared on BHI + CM plates after 24-48 hr incubation at 30°C was picked and homogenized in PBS, then serially diluted up to $10^{-6}$ before spreading on BHI agar + CM + X-gal. Blue colonies indicated plasmid integration. Several blue colonies were examined using colony PCR to determine the side of plasmid integration. To excise the plasmids, a selection of these colonies (representing both sides of integration, if possible) was then inoculated individually into 10 ml of BHI broth and incubated with shaking until saturation. We serially diluted the saturated broth and plated out the $10^{-4}$-$10^{-6}$ dilutions onto BHI agar + Xgal and incubated the plates overnight at 37°C. White colonies were picked and reinoculated on fresh BHI + Xgal plates as well as BHI + CM and incubated overnight at 37°C. Colonies which remained white on BHI + Xgal and were unable to grow on BHI + CM were examined by colony PCR to determine whether the recombination template had been successfully integrated. Colonies with the expected amplicon size (indicating KO of target gene) were kept. Genomic DNA was isolated from these colonies and sent for WGS (Azenta).
